# FGF2 disruption enhances thermogenesis in brown and beige fat to protect against obesity and hepatic steatosis

**DOI:** 10.1101/2020.12.02.407650

**Authors:** Haifang Li, Xinzhi Zhang, Cheng Huang, Huan Liu, Shuang Liu, Qiang Zhang, Mei Dong, Mengjie Hou, Yiming Liu, Hai Lin

**Author notes:** Corresponding authors: Haifang Li, Hai Lin. **Author contributions:** Haifang Li, Xinzhi Zhang, Cheng Huang, and Huan Liu performed in *vitro* and in *vivo* experiments, wrote the manuscript, organized the literature and figures and performed statistical analysis. Haifang Li, and Hai Lin conceived the project, led and supervised the study, and reviewed/edited the manuscript. Shuang Liu, Qiang Zhang, Mei Dong, Mengjie Hou, and Yiming Liu performed experiments and contributed to discussion.

## Abstract

Since brown and beige fat expend energy in the form of heat via non-shivering thermogenesis, identifying key regulators of thermogenic functions represents a major goal for development of potential therapeutic avenues for obesity and associated disorders. Here, we identified fibroblast growth factor 2 (FGF2) as a novel thermogenic regulator. FGF2 gene disruption resulted in increased thermogenic capability in both brown and beige fat, which was supported by increased UCP1 expression, enhanced respiratory exchange ratio, and elevated thermogenic potential under cold challenge or β-adrenergic stimulation. Thus, deletion of FGF2 protected mice from high fat-induced obesity and hepatic steatosis. Mechanistically, FGF2 acts in autocrine/paracrine fashions *in vitro*. Exogenous FGF2 supplementation inhibits both PGC-1α and PPARγ expression through ERK phosphorylation, thereby limiting PGC-1α/PPARγ interactions, and leading to suppression of UCP1 expression and thermogenic activity in brown and beige adipocytes. These findings suggest a viable potential strategy for use of FGF2-selective inhibitors in treatment of combating obesity and related disorders.

## INTRODUCTION

In mammals, there are three types of adipose tissue: white, brown, and beige (*Rosen and Spiegelman, 2014*). While white adipose tissue (WAT) serves as a repository for fatty acids, brown and beige adipocytes burn fatty acids and glucose to generate heat, leading to increased energy expenditure (*Rosen and Spiegelman, 2014*; *Poekes et al., 2015*; *Wu et al., 2012*). Brown adipocytes derive from a myf5-positive cell lineage, characterized by the presence of small, dense lipid droplets enriched with mitochondria and high expression of uncoupling protein 1 (UCP1), which enables the uncoupling of oxidative phosphorylation from ATP production (*Poekes et al., 2015*; *Stanford et al., 2013*). The expression of UCP1 is driven by peroxisome proliferator-activated receptor gamma (PPARγ) in cooperation with other transcriptional components, including PPARγ coactivator-1α (PGC-1α) (*Wu et al., 1999*; *Puigserver and Spiegelman, 2003*). In contrast, beige adipocytes emerge from white fat through a process called browning or beiging (*Wu et al., 2012*). The PGC-1α transcriptional cofactor is also critical important to control white-to-beige fat conversion (*Puigserver and Spiegelman, 2003*; *Xue et al., 2005*). Similar to brown adipocytes, beige adipocytes possess, albeit to a lesser degree, several features indispensable to thermogenesis, such as multilocular lipid droplet morphology, high UCP1 expression, and densely packed mitochondria (*Rosen and Spiegelman, 2014*; *Wu et al., 2012*). Previous studies have provided evidence that metabolically active brown and beige fat are present in adult humans (*Virtanen et al., 2009*; *Rogers, 2015*), and their abundance is inversely correlated with fat mass and obesity-associated disorders (*Virtanen et al., 2009*; *Saito et al., 2009*).

Large scale studies have demonstrated the inducibility of both brown and beige fat (*van Marken Lichtenbelt et al., 2009*; *Seale et al., 2011*; *Bachman et al., 2002*; *Villarroya and Vidal-Puig, 2013*). β-adrenergic signaling induced by cold exposure and/or β-adrenergic agonists undoubtedly serve as the primary physiological signal pathway for activation of brown fat thermogenesis and stimulation of beige adipocyte development (*van Marken Lichtenbelt et al., 2009*; *Bachman et al., 2002*). In addition, several other secreted factors and hormones, such as BMP8 (*Whittle et al., 2012*), FGF21 (*Fisher et al., 2012*), Irisin (*Boström et al., 2012*), and apelin (*Than et al., 2015*), have been shown to participate in regulating thermogenic activity in brown and/or beige adipocytes, the activation of which may profoundly decrease fat accumulation while improving lipid metabolism and glucose homeostasis. Thus, identifying key regulators of the thermogenic functions of brown and beige adipocytes represents a major goal for development of potential therapeutic avenues for obesity and metabolic diseases, such as fatty liver and type II diabetes (*Villarroya and Vidal-Puig, 2013*; *Zeve et al., 2009*).

Fibroblast growth factor 2 (FGF2), also known as basic FGF (bFGF), is among the first recognized members of the FGF family (*Powers et al., 2000*). Through loss-of-function studies, FGF2 has been reported to play essential or predominant roles in the development of vessels (*Zhou et al., 1998*), nerves (*Raballo et al., 2000*), and bone (*Hurley et al., 1998*; *Montero et al., 2000*). Additionally, the regulation of white adipogenic differentiation by FGF2 has been rigorously demonstrated (*Kawaguchi et al., 1998*; *Kakudo et al., 2007*; *Hao et al., 2016*; *Xiao et al., 2010*). Kawaguchi *et al*. showed the induction of *de novo* adipogenesis in reconstituted basement membrane supplemented with FGF2 (*Kawaguchi et al., 1998*). FGF2 significantly enhances the adipogenic differentiation of human adipose-derived stem cells (hASCs) and the expression of PPARγ (*Kakudo et al., 2007*). Hao *et al*. described a positive correlation between plasma FGF2 levels and fat mass, as well as an increased risk of obesity (*Hao et al., 2016*). However, work by Xiao *et al*. reported that bone marrow stem cells of FGF2-deficient mice showed enhanced lipogenic ability with up-regulation of key adipogenic signaling molecules (*Xiao et al., 2010*).

Given that FGF2 is correlated with white adipogenesis and fat mass, we hypothesized that it may participate in regulating the thermogenic function of brown and/or beige fat. Here, we unexpectedly discovered that FGF2 gene disruption strongly enhanced the thermogenic action of both brown and beige fat, which led to an increase in energy expenditure and improvement of lipid homeostasis. Consequently, FGF2 gene knockout (KO) alleviated high fat diet (HFD)-induced obesity and hepatic steatosis. Mechanistically, FGF2 suppression of PGC-1α and PPARγ expression and interaction led to attenuated UCP1 expression and thermogenic activity in brown and beige adipocytes, which was partially mediated by the ERK signaling. These findings established that FGF2 negatively regulates thermogenesis in both brown and beige fat, thus suggesting a strong potential therapeutic approach for the treatment of obesity-associated metabolic disorders via FGF2-specific signaling inhibitors.

## RESULTS

### FGF2 gene disruption is associated with enhanced brown fat thermogenesis and beiging of white fat

To evaluate the role of FGF2 in the thermogenic potential of adipose tissues, we generated FGF2-KO mice (genetic background C57BL/6J) (*figure supplement 1*). To initially characterize the KO phenotype, we fed 3-week-old male FGF2^+/+^ and FGF2^−/−^ mice with chow diet for 14 weeks and found that although the dynamic changes in body weight and average body weight were indistinguishable between groups at 17 weeks old (*figure supplement 2A,B*), FGF2^−/−^ mice consumed more food than WT littermates during the course of the experiment (*figure supplement 2C*). Correspondingly, FGF2^−/−^ mice had a significantly higher whole feed/gain ratio (*figure supplement 2D*). Notably, the interscapular BAT (iBAT) index of FGF2^−/−^ mice was significantly lower than that of FGF2^+/+^ mice (*Figure 1A,B-figure supplement 2E*). However, in contrast to control, FGF2^−/−^ mice had markedly less subcutaneous fat (*figure supplement 2E*). Specifically, the inguinal WAT (iWAT) index was substantially lower (p=0.057) in FGF2^−/−^ mice than those in FGF2^+/+^ mice (*Figure 1D*), clearly indicated by the representative graphs of iWAT (*Figure 1C*).

**Fig. 1.**
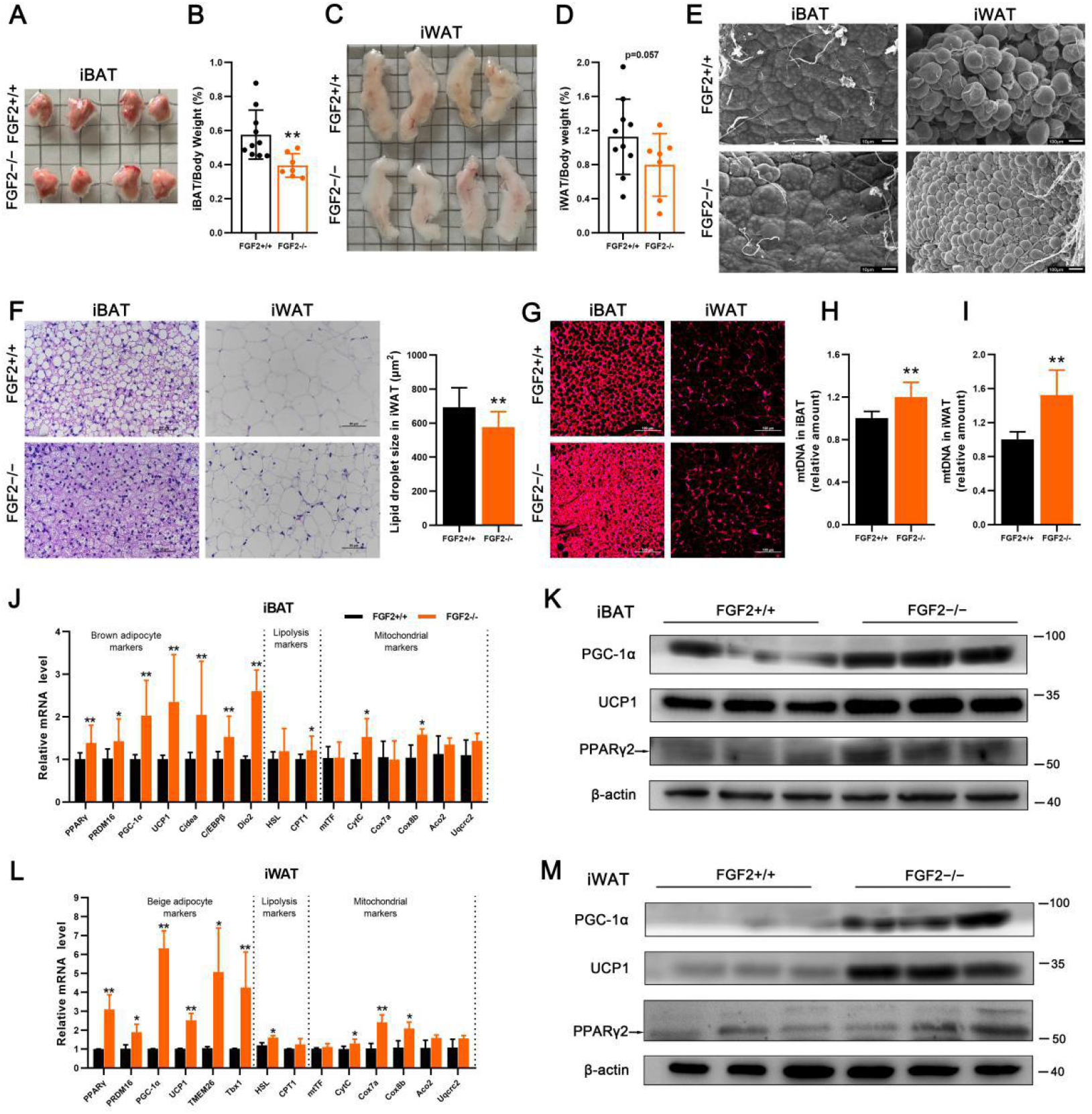
FGF2 gene disruption leads to enhanced brown fat thermogenesis and beiging of white fat. Tissue samples were collected from 17-week-old FGF2^+/+^ and FGF2^−/−^ mice. (*A* and *B*) Representative iBAT images and ratios of iBAT/body weight of FGF2^+/+^ and FGF2^−/−^ mice. (*C* and *D*) Representative iWAT images and ratios of iWAT/body weight of FGF2^+/+^ and FGF2^−/−^ mice. Values represent means ± SEM (n = 7∼10). \*\**p*<0.01 compared with FGF2^+/+^ mice. (*E*) Representative scanning electron microscopy images of iBAT (Scale bar = 10 μm) and iWAT (Scale bar = 100 μm) sections. (*F*) Representative images of H&E staining of iBAT and iWAT sections, and the lipid droplet size in iWAT. Scale bar = 50 μm. (*G*) Immunofluorescence staining of UCP1 (red) in FGF2^+/+^ and FGF2^−/−^ iBAT and iWAT sections. Scale bar = 100 μm. (*H* and *I*) Quantification of relative mtDNA levels in iBAT (*H*) and iWAT (*I*) in FGF2^+/+^ and FGF2^−/−^ mice. (*J*) The relative mRNA levels of brown adipocyte, lipolysis, and mitochondrial markers in iBAT of FGF2^+/+^ and FGF2^−/−^ mice, determined by qRT-PCR. (*K* and *M*) Western blot analysis of PPARγ, PGC-1α, and UCP1 protein contents in iBAT (*K*) and iWAT (*M*) of FGF2^+/+^ and FGF2^−/−^ mice. Blots were stripped and then probed with β-actin to normalize for variation in loading and transfer of proteins. (*L*) The relative mRNA levels of beige adipocyte, lipolysis, and mitochondrial markers in iWAT of FGF2^+/+^ and FGF2^−/−^ mice, determined by qRT-PCR. Values represent means ± SEM (n = 6). \**p*<0.05, \*\**p*<0.01 compared with FGF2^+/+^ samples.

Histologically, iBAT in KO was comparable with that in WT tissue (*Figure 1E,F*). However, the UCP1 immuno-reactivity and mtDNA levels in KO iBAT were evidently elevated (*Figure 1G,H*). Interestingly, the transcription levels of thermogenic-associated genes (PPARγ, PRDM16, PGC-1α, UCP1, Cidea, C/EBPβ, and Dio2) and mitochondrial markers (CytC, and Cox8b) were drastically increased in FGF2^−/−^ iBAT comparing with those in the WT iBAT (*Figure 1J*). Meanwhile, CPT1, a gene associated with fatty acid oxidation, was also transcriptionally activated in iBAT of FGF2^−/−^ mice (*Figure 1J*). In addition, the protein levels of PGC-1α, UCP1, and PPARγ were all elevated in iBAT of FGF2^−/−^ mice (*Figure 1K*). Notably, the adipogenic potential of KO iBAT was enhanced, as the mRNA levels of aP2, FAS and LPL, and the protein expression of aP2 were all highly induced (*figure supplement 3A,C*). These data indicate that FGF2-KO mice recruit and expend more fat in BAT via elevated thermogenesis.

Moreover, FGF2 disruption led to a marked decrease in lipid droplet size in iWAT (*Figure 1E,F*), characteristic of reduced triglyceride accumulation or accelerated triglyceride release. Strikingly, immunofluorescence analysis revealed greater intensity of UCP1 immuno-reactivity in the iWAT of FGF2-KO mice (*Figure 1G*). Notably, the mtDNA copy number in KO iWAT was elevated by ∼50% compared with that in the WT tissue (p<0.01) (*Figure 1I*). Furthermore, iWAT from FGF2^−/−^ mice displayed elevated transcription of thermogenic markers (PPARγ, PRDM16, PGC-1α and UCP1), beige adipocyte-specific genes (TMEM26 and Tbx1), and mitochondrial markers (CytC, Cox7a, and Cox8b) (*Figure 1L*). Likewise, KO iWAT also showed increased levels of thermogenic marker proteins, *e*.*g*., PPARγ, PGC-1α and UCP1 (*Figure 1M*). Moreover, the transcription of adipogenic-related genes (C/EBPα, FAS, LPL, and aP2) and the protein expression of aP2 were induced in KO iWAT (*figure supplement 3B,D*). These results suggest that the decrease in adipocyte size in FGF2^−/−^ iWAT depends greatly on the degree of induced beiging and the thermogenic potential rather than on reduced triglyceride synthesis.

### FGF2-KO mice show increased respiratory exchange ratio (RER) and body temperature, as well as activated responses to cold challenge and β3-AR stimulation

In view of the role of brown and beige fat in non-shivering thermogenesis, we compared the respiratory metabolic parameters of the FGF2^+/+^ and FGF2^−/−^ mice at ambient temperature (25°C). In contrast to WT mice, carbon dioxide production and RER in FGF2^-/-^ mice both increased substantially in the dark (p<0.05), and oxygen consumption was also markedly elevated (p=0.075) (*Figure 2A-C*). In the light, oxygen consumption, carbon dioxide production, and RER were also higher in KO mice, although not significantly (*Figure 2A-C*).

**Fig. 2.**
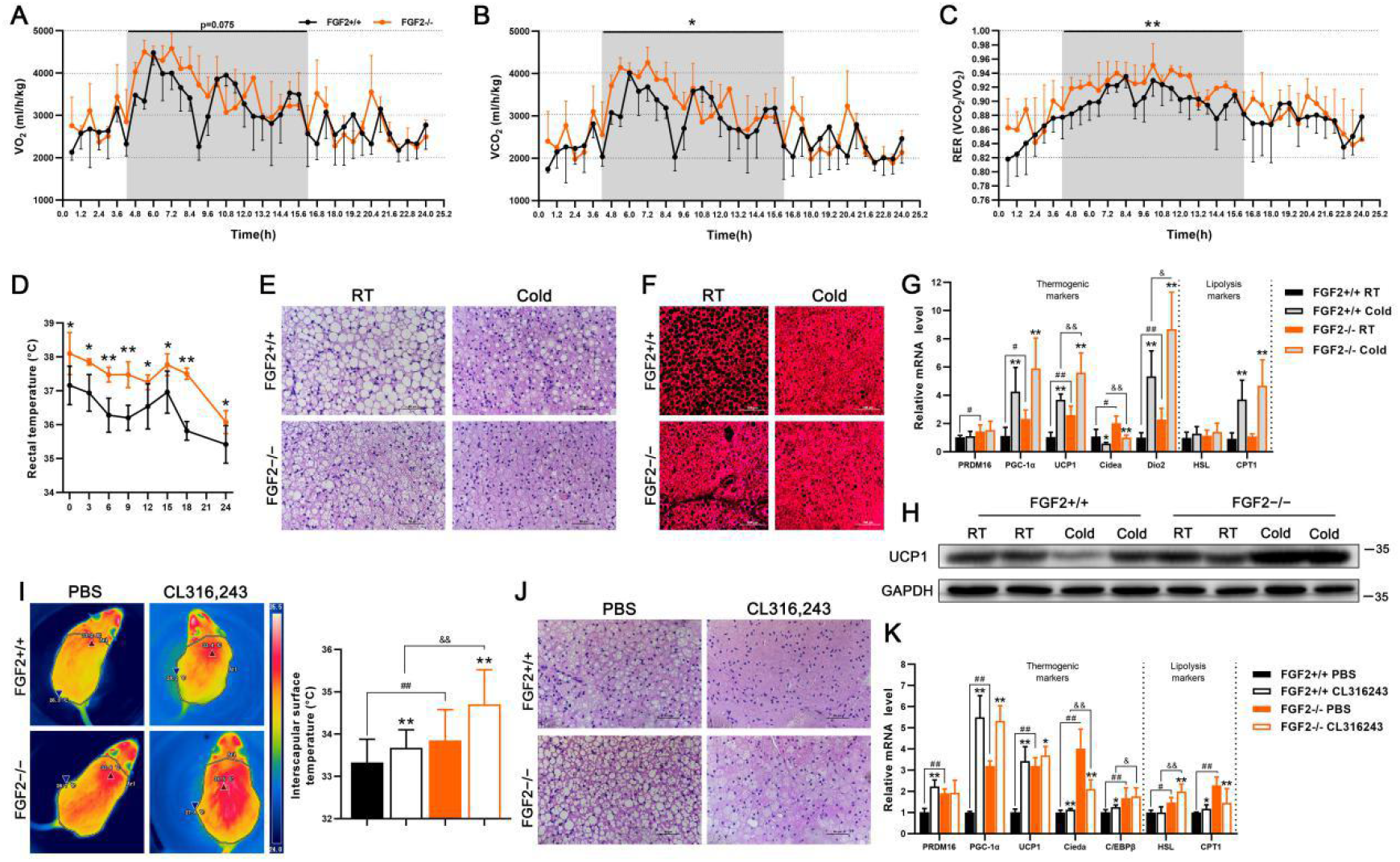
FGF2-KO mice show increased whole-body energy expenditure and activated thermogenic capability of brown fat. (*A* and *B*) O_2_ consumption (*A*) and CO_2_ production (*B*) (expressed as ml/h/kg) measured in 12-week-old FGF2^+/+^ and FGF2^−/−^ mice during a 24-hour period measured using a CLAMS apparatus. (*C*) RER dynamics calculated by VCO_2_/VO_2_. (*D*) Core body temperature changes in FGF2^+/+^ and FGF2^−/−^ mice following cold challenge, determined by rectal probe every 3 or 6 hours for a 24-h duration. (*E* and *F*) Representative images of H&E staining (Scale bar = 50 μm) (*E*) and immunofluorescence staining of UCP1 (red) (Scale bar = 100 μm) (*F*) of iBAT sections, under room temperature (RT) or cold challenge for 24 h. (*G*) qRT-PCR analysis of thermogenic- and lipolysis-related gene expression in FGF2^+/+^ and FGF2^−/−^ iBAT under normal temperature or cold challenge for 24 h. (*H*) Western blot analysis of UCP1 protein levels in FGF2^+/+^ and FGF2^−/−^ iBAT under RT or cold challenge for 24 h. GAPDH was used as a loading control. (*I*) Representative thermal images and dorsal interscapular surface temperatures of FGF2^+/+^ and FGF2^−/−^ mice after injection with CL316,243 or PBS control. (*J*) Representative images of H&E staining of iBAT sections upon PBS or CL316,243 injection. Scale bar = 50 μm. (*K*) qRT-PCR analysis of thermogenic- and lipolysis-related gene expression in FGF2^+/+^ and FGF2^−/−^ iBAT under PBS or CL316,243 treatments. Data represent means ± SEM. \**p*<0.05, \*\**p*<0.01 vs. Vehicle in the same littermates; ^#^*p*<0.05, ^##^*p*<0.01 vs. FGF2^+/+^ mice upon Vehicle treatment; ^&^*p*<0.05, ^&&^*p*<0.01 vs. FGF2^+/+^ mice upon cold challenge or CL316,243 treatment.

Given that increased metabolic rate is often accompanied by higher thermogenic capacity, we examined the animals’ body temperature under cold challenge conditions. As expected, FGF2^-/-^ mice showed elevated rectal temperature both prior to and during the 24-hour cold challenge, which was approximately 1°C higher on average than that of WT littermates at corresponding time points (*Figure 2D*). By using an infrared camera, we further found that the surface temperature of KO iBAT was significantly higher than that of WT, in both untreated and CL316,243 (CL) treatments (*Figure 2I*).

In light of our findings of potentiated activation of body temperature in KO mice under both cold and β3-AR stimulation conditions, we further examined the thermogenic markers in treated adipose tissues. In iBAT and iWAT, both cold challenge and β3-AR stimulation led to the occurrence of smaller lipid droplets, and higher expression of thermogenic genes (*Figure 2E-H,J,K, Figure 3*). However, the eWAT adipocytes were unaffected by CL treatment (*figure supplement 4*), consistent with previous research that indicated brown-like fat cells are seldom observed in epididymal/perigonadal adipose tissue even under cold challenge or β3-AR stimulation (*Seale et al., 2011*). Notably, in response to cold challenge, FGF2^−/−^ iBAT exhibited a strongly potentiated induction of UCP1, and Dio2 mRNA expression, as well as an elevated UCP1 protein level relative to WT controls (*Figure 2G,H*). However, following CL injection, the mRNA levels of Cidea, C/EBPβ, and HSL were significantly higher in FGF2^−/−^ iBAT than those in FGF2^+/+^ iBAT (*Figure 2K*). Moreover, KO iWAT showed a more pronounced increase of PRDM16, PGC-1α, UCP1, TMEM26, Tbx1, and CPT1 transcription levels, and a higher UCP1 protein content upon cold challenge (*Figure 2C,D*), and only an elevated transcription of Tbx1 following CL treatment (*Figure 2F*) compared with WT iWAT. These data reflect a highly stronger cold-response and relatively a slight higher CL-responsiveness by both KO iBAT and iWAT than by WT tissues.

**Fig. 3.**
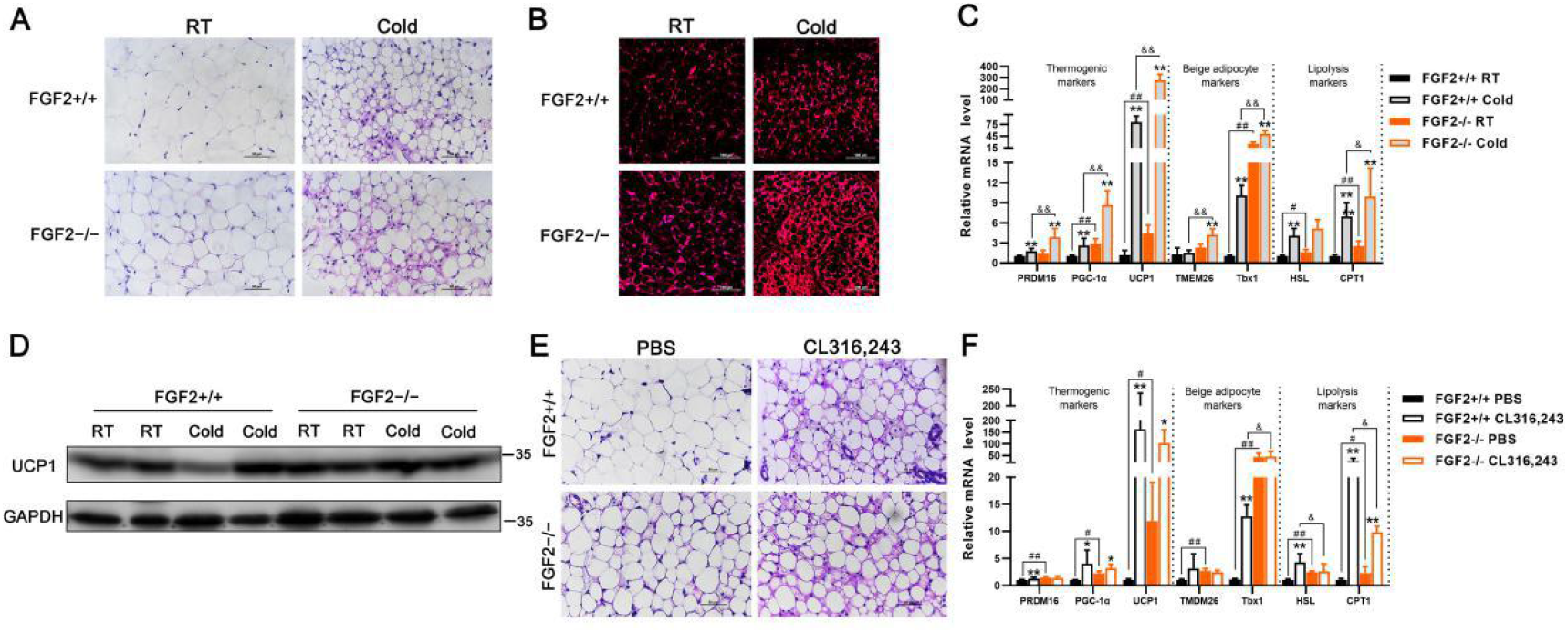
The thermogenic capability of iWAT exhibits higher potentiation in FGF2-KO mice than in WT, under either cold challenge or β3-AR stimulation conditions. (*A* and *B*) Representative images of H&E staining (Scale bar = 50 μm) (*A*) and immunofluorescence staining of UCP1 (red) (Scale bar = 100 μm) (*B*) of iWAT sections under RT or cold challenge for 24 h. (*C* and *F*) qRT-PCR analysis of thermogenic-, beige adipocyte-, and lipolysis-related gene expression in FGF2^+/+^ and FGF2^−/−^ iWAT under normal temperature or cold challenge for 24 h (C), or following PBS or CL316,243 treatments (*F*). (*D*) Western blot analysis of UCP1 protein levels in FGF2^+/+^ and FGF2^−/−^ iBAT under RT or cold challenge for 24 h. GAPDH was used as a loading control. (*E*) Representative images of H&E staining of iWAT sections following PBS or CL316,243 treatments. Scale bar = 50 μm. Data represent means ± SEM. \**p*<0.05, \*\**p*<0.01 vs. Vehicle in the same littermates; ^#^*p*<0.05, ^##^*p*<0.01 vs. FGF2^+/+^ mice upon Vehicle treatment; ^&^*p*<0.05, ^&&^*p*<0.01 vs. FGF2^+/+^ mice upon cold challenge or CL316,243 treatment.

### FGF2-KO mice show higher stability in lipid homeostasis and amelioration of HFD-induced obesity and hepatic steatosis

In view of the enhanced function of brown and beige fat resulting from FGF2*-*KO, its effects on lipid homeostasis were next investigated. Upon chow feeding, significantly reduced plasma TG but not of TCH content was observed in the FGF2-disrupted mice (*figure supplement 5A,B*). The ALT and AST activities were also substantially decreased by FGF2 deficiency (p=0.089 for ALT, p<0.01 for AST) (*figure supplement 5C,D*). In response to HFD feeding, the plasma TG content was significantly lower in FGF2-KO mice than in WT animals (p<0.05), while TCH content, ALT and AST activities were modestly down-regulated (p>0.05) (*figure supplement 5A-D*).

HFD is prone to induce obesity and ectopic fat deposition in livers. Therefore, we determined the influence of FGF2 disruption on fat accumulation and hepatic steatosis following 14 weeks of HFD feeding. Interestingly, compared with WT, HFD-fed FGF2-KO mice exhibited significantly lower body weight, with an elevated feed/gain ratio (*figure supplement 6A,B*). Specifically, dual energy X-ray absorptiometry (DXA) and anatomical imaging revealed vastly lower subcutaneous fat mass in FGF2-KO mice, compared with that in WT (*Figure 4A*-*figure supplement 6C*). Moreover, the iWAT index was greatly down-regulated, while the iBAT index was not significantly altered in HFD-fed KO mice (*Figure 4B-E*). In addition, the adipocyte size in iWAT and lipid droplet size in iBAT were generally smaller among FGF2-KO mice than among WT (*Figure 4F*).

**Fig. 4.**
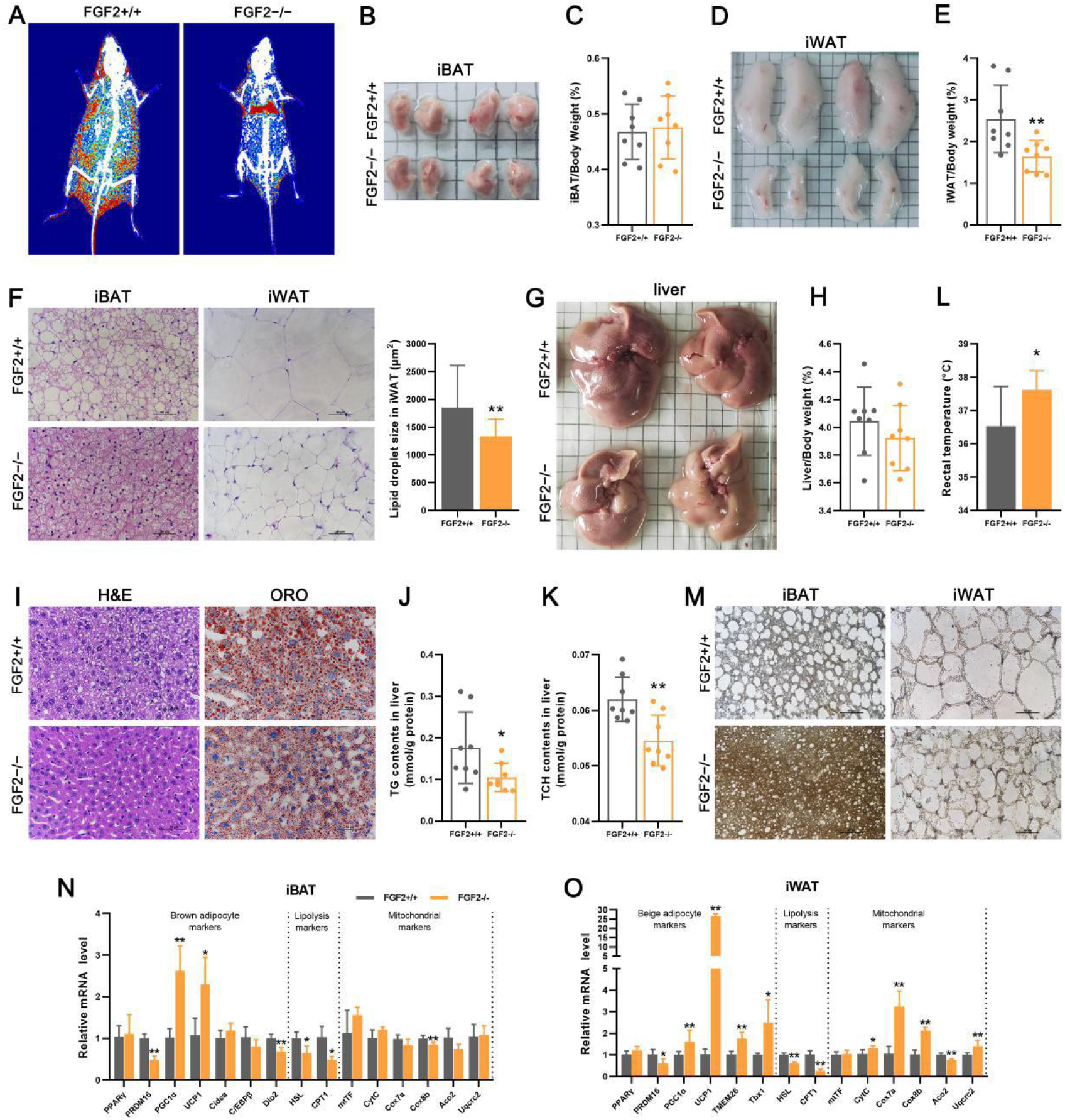
FGF2-KO protects mice against HFD-induced obesity and hepatic steatosis, primarily due to activated thermogenic function in both brown and white fats. (A) Representative DXA images of 14-week-old FGF2^+/+^ and FGF2^−/−^ mice fed with HFD. Red represents areas with more than 50% fat. (*B* and *C*) Representative iBAT images and ratios of iBAT/body weight of HFD-fed FGF2^+/+^ and FGF2^−/−^ mice at age of 17 weeks old. (*D* and *E*) Representative iWAT images and ratios of iWAT/body weight of HFD-fed FGF2^+/+^ and FGF2^−/−^ mice at age of 17 weeks old. (*F*) Representative images of H&E staining of iBAT and iWAT sections, and the lipid droplet size in iWAT. Scale bar = 50 μm. (*G* and *H*) Representative liver images (*G*) and the liver/body weight ratios (*H*) of FGF2^+/+^ and FGF2^−/−^ HFD-fed mice. (*I*) Representative images of H&E and ORO staining of liver sections. Scale bar = 50 μm. (J and *K*) TG (*J*) and TCH (*K*) contents in the livers of HFD-fed FGF2^+/+^ and FGF2^−/−^ mice. (*L*) The core body temperature of HFD-fed FGF2^+/+^ and FGF2^−/−^ mice. (*M*) Immunohistochemistry staining of UCP1 (brown) in HFD-fed FGF2^+/+^ and FGF2^−/−^ iBAT and iWAT sections. Scale bar = 50 μm. (*N* and *O*) The relative mRNA levels of brown/beige adipocyte-, lipolysis-, and mitochondrial-related markers in iBAT (*N*) and iWAT (*O*) of HFD-fed FGF2^+/+^ and FGF2^−/−^ mice, determined by qRT-PCR. Data represent means ± SEM (n = 8).\**p*<0.05, \*\**p*<0.01 vs. HFD-fed FGF2^+/+^ mice.

Strikingly, the livers of FGF2^−/−^ mice were visibly smaller and the liver index was slight lower than WT fed with HFD (*Figure 4G,H*), thus indicating that the HFD-induced fatty liver phenotype may be ameliorated by FGF2 disruption. As expected, H&E and ORO staining of liver sections (*Figure 4I*), as well as lower hepatic levels of TG and TCH (*Figure 4J,K*), gave further evidence that FGF2 KO led to alleviation of HFD-induced hepatic steatosis. Furthermore, liver-specific expression of fat synthesis- or fatty acid oxidation-associated genes were not significantly different between HFD-fed KO mice and WT, except for a decrease of ACC (*figure supplement 7*), suggesting that the amelioration of hepatic fat deposition was not attributable to the direct alteration of *in situ* fat metabolism.

To ascertain whether the amelioration of HFD-induced obesity and hepatic steatosis mediated by FGF2 deficiency was due to the elevation of thermogenic activity, we compared core body temperature and thermogenic gene expression between HFD-fed WT and KO mice. We found that rectal temperature was significantly higher in HFD-fed KO mice compared to that in WT (*Figure 4L*). Moreover, HFD-fed KO mice displayed substantially greater UCP1 immunostaining in both iBAT and iWAT sections (*Figure 4M*). In addition, the transcriptional levels of PGC-1α and UCP1 in iBAT, as well as those of PGC-1α, UCP1, TMEM26, Tbx1, CytC, Cox7a, Cox8b, and Uqcrc2 in iWAT, were all profoundly elevated in HFD-fed KO mice (*Figure 4N,O*). These results suggested that the alleviated HFD-induced obesity and hepatic steatosis phenotype associated with FGF2 deficiency is due in large measure to the enhanced thermogenic ability of brown and beige fats.

### FGF2-KO-derived brown and beige adipocytes exhibit higher thermogenic gene expression *in vitro*

Given the enhancement of *in vivo* thermogenesis resulting from FGF2 disruption, *in vitro* experiments were conducted to compare the thermogenic gene expression levels between WT- and KO-derived adipocytes. We found that the differentiated FGF2^−/−^ brown adipocytes exhibited higher thermogenic gene transcription, including PPARγ, PGC-1α, UCP1, and C/EBPβ, as well as lipolysis markers (HSL and CPT1) and the mitochondrion marker mtTF than did controls (*Figure 5A*). Similarly, the protein expression of PPARγ, PGC-1α, and UCP1 was enhanced in FGF2^−/−^ brown adipocytes, compared to that in WT cells (*Figure 5B*). In addition, the mitochondria density and the UCP1 immuno-reactivity were also elevated in KO brown adipocytes (*Figure 5C,D*). We used isoproterenol (ISO) to induce the beiging of *in vitro* cultured WT- and KO-derived white adipocytes, and identified that the beiging-associated genes, including PPARγ, PGC-1α, UCP-1, and TMEM26 were all activated in KO cells (*Figure 5E*). Moreover, in contrast to WT, the KO beiged white adipocytes displayed enhanced PGC-1α and UCP1 protein levels, mitochondria density, and UCP1 immuno-reactivity (*Figure 5F-H*). These results indicated that FGF2-KO brown and beige adipocytes also show higher thermogenic gene expression *in vitro*, hinting the antocrine regulation of FGF2 on thermogenesis.

**Fig. 5.**
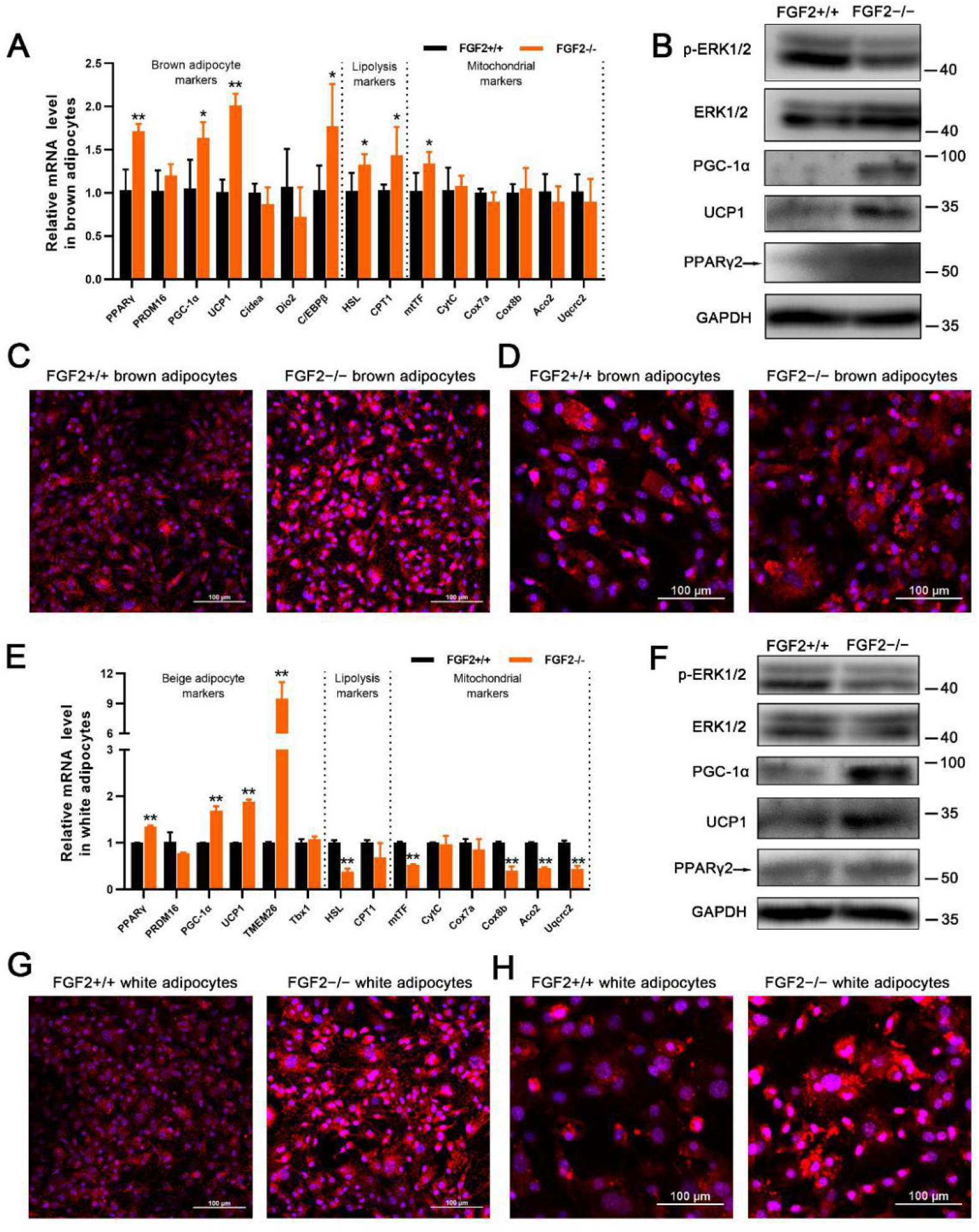
FGF2^−/−^-derived brown and beige adipocytes exhibit higher thermogenic gene expression *in vitro*. (*A*) Relative mRNA levels of brown adipocyte, lipolysis, and mitochondrial genes in differentiated FGF2^+/+^ and FGF2^−/−^ brown adipocytes. (*B*) Western blot analysis of PPARγ, PGC-1α, UCP1, p-ERK, and ERK protein contents in brown adipocytes derived from FGF2^+/+^ and FGF2^−/−^ mice. GAPDH was used as a loading control. (*C* and *D*) MitoTracker staining (red) (*C*) and immunofluorescence staining of UCP1 (red) (*D*) in brown adipocytes derived from WT and KO mice. The nuclei (blue) were stained with DAPI. Scale bar = 100 μm. (*E*) Relative mRNA expression beige adipocyte, lipolysis, and mitochondrial genes in differentiated FGF2^+/+^ and FGF2^−/−^ white adipocytes. (*F*) Western blot analysis of PPARγ, PGC-1α, UCP-1, p-ERK, and ERK protein contents in beiging white adipocytes derived from FGF2^+/+^ and FGF2^−/−^ mice. GAPDH was used as a loading control. (*G* and *H*) MitoTracker staining (red) (*G*) and immunofluorescence staining of UCP1 (red) (*H*) of beiging white adipocytes derived from WT and KO mice. The nuclei (blue) were stained with DAPI. Scale bar = 100 μm. Data represent means ± SEM (n = 6). \**p*< 0.05, \*\**p*< 0.01 compared with that from FGF2^+/+^ mice.

### Exogenous FGF2 application inhibits expression of thermogenic genes in cultured brown and white adipocytes, partially through activating ERK phosphorylation

Previous reports have established that FGF2 acts in either an autocrine or a paracrine fashion via FGFR1 binding that requires concurrent interaction with heparin (HP) (*Zhou et al., 1998*). Thus, we supplemented exogenous FGF2 together with HP to the cell culture medium to test the alteration of thermogenic genes. In the induced brown adipocytes, FGF2 plus HP supplementation (10 ng/mL each) strongly suppressed the mRNA levels of PGC-1α, UCP1, and CytC, as well as the protein expression of PGC-1α (*figure supplement 8*). Thus, we used FGF2 in conjunction with HP in all the following experiments. Notably, the FGFR1 inhibitor SSR128129E (SSR) was able to antagonize the inhibition by FGF2 on thermogenic gene expression (*figure supplement 9*). This supported the paracrine regulation of FGF2 on thermogenesis of brown adipocytes.

As a growth factor, FGF2 participates in activation of some growth-related signals, such as the ERK signaling pathway (*Kim et al., 2015*). We observed a decrease of p-ERK/ERK ratio in both cultured FGF2^−/−^ brown and beige adipocytes (*Figure 5B,F*). Herein, two-day-induced WT brown adipocytes were treated with exogenous FGF2 over a one-hour time course to further test the involvement of ERK signaling. We can see that exogenous FGF2 application led to a rapid induction of ERK1/2 phosphorylation (*Figure 6A*). Furthermore, an ERK-specific inhibitor PD0325901 (PD) was included in the differentiation medium for WT brown adipocytes in the absence or presence of FGF2. Interestingly, FGF2 transcriptional suppression of thermogenic markers was substantially decreased by PD application (*Figure 6B*). Moreover, PD also negated the FGF2-mediated blockade of PGC-1α, UCP1, and PPARγ protein expression (*Figure 6C*). These data clarify the involvement of ERK signaling in the FGF2-mediated transcriptional and translational blockade of thermogenic genes.

**Fig. 6.**
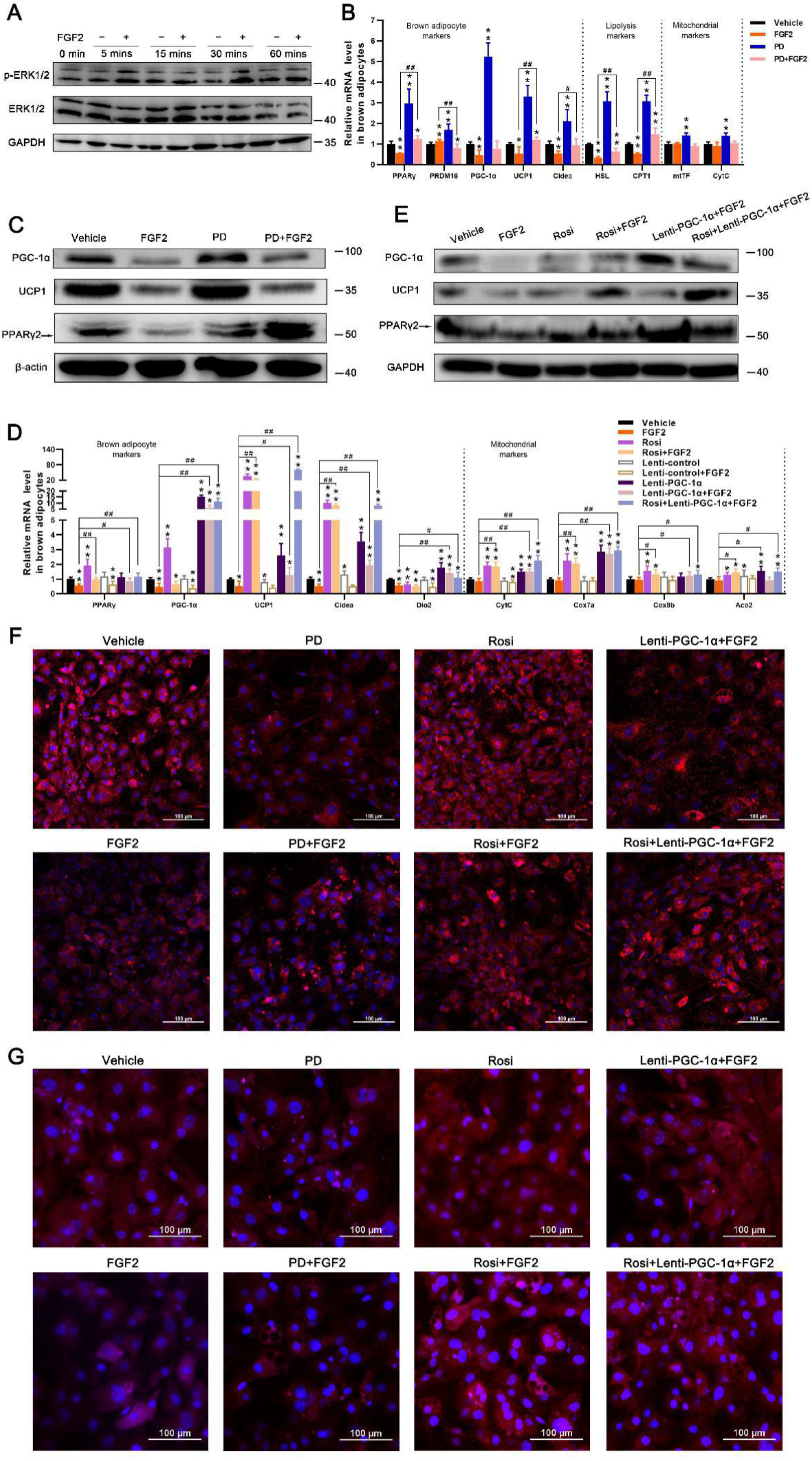
FGF2 inhibits thermogenic gene expression in brown adipocytes *in vitro*, via ERK signaling-induced PPARγ and PGC-1α suppression. (*A*) Expression of ERK1/2 and p-ERK1/2 proteins in differentiated brown SVFs after supplementation with FGF2 or Vehicle for 5, 15, 30, and 60 min, determined by western blotting. Blots were stripped and probed again with GAPDH to normalize for variation in loading and transfer of proteins. (*B*) Relative mRNA levels of brown adipocyte-, lipolysis-, and mitochondrial-associated genes in Vehicle, FGF2, PD, or PD+FGF2 -treated brown adipocytes, determined by qRT-PCR. GAPDH serves as a loading control. (*C*) Protein expression of PPARγ, PGC-1α, and UCP1 in cells treated as in (*B*), determined by western blotting. β-actin serves as a loading control. (*D*) Relative mRNA levels of brown adipocyte-, lipolysis-, and mitochondrial-associated genes in Vehicle, FGF2, Rosi, Rosi+FGF2, Lenti-control, Lenti-control+FGF2, Lenti-PGC-1α, Lenti-PGC-1α+FGF2, or Rosi+Lenti-PGC-1α+FGF2 -treated brown adipocytes, determined by qRT-PCR. (*E*) Protein expression of PPARγ, PGC-1α, and UCP1 in brown adipocytes treated with Vehicle, FGF2, Rosi, Rosi+FGF2, Lenti-PGC-1α+FGF2, or Rosi+Lenti-PGC-1α+FGF2, determined by western blotting. (*F* and *G*) MitoTracker staining (red) (*F*) and immunofluorescence staining of UCP1 (red) (*G*) of treated brown adipocytes. The nuclei (blue) were stained with DAPI. Scale bar = 100 μm. Data represent means ± SEM.\**p*<0.05, \*\**p*<0.01 vs. Vehicle; ^#^*p*<0.05, ^##^*p*<0.01 vs. FGF2 treatment.

Strikingly, FGF2 also robustly inhibited the expression of beiging-related genes in ISO-induced beiging white adipocytes, but were transcriptionally restored by application of the FGFR1 inhibitor SSR (*figure supplement 10*). These results demonstrate that FGF2 also plays a negative regulatory role in the beiging of white adipocytes through a paracrine-dependent manner. Besides, the activated ERK signaling contributed to negative regulation of beiging of white adipocytes by FGF2 (*figure supplement 11A-C*).

Together, these data illustrate that exogenous FGF2 application is able to inhibit thermogenic gene expression in both brown and beige adipocytes in paracrine fashions, which at least partially via activation of ERK1/2 phosphorylation.

### PPARγ and PGC-1α cooperatively participate in FGF2 suppression of thermogenic gene expression in brown and white adipocytes

The PPARγ transcription factor controls thermogenic gene expression in conjunction with other regulatory co-factors, such as PGC-1α (*Puigserver and Spiegelman, 2003*; *Ahmadian et al., 2013*). Notably, accompanied by induction of UCP1 expression, PPARγ and PGC-1α mRNA and protein levels were also elevated by FGF2 deficiency in both iBAT and iWAT (*Figure 1J*-*M*). Similarly, in *in vitro* cultures of brown and beiging white adipocytes, FGF2 substantially suppressed PPARγ and PGC-1α accumulation (*Figure 6B,C*-*figure supplement 11B,C*). We therefore employed rosiglitazone (Rosi), a PPARγ-specific agonist, and a recombinant PGC-1α expression lentivirus construct (Lenti-PGC-1α) to test whether PPARγ and PGC-1α performed essential functions in the FGF2-mediated pathway for suppression of thermogenic gene expression.

In differentiated brown adipocytes, we found that Rosi counteracted FGF2 inhibition of UCP1 and Cidea transcription, and Lenti-PGC-1α also restored UCP1, Cidea, and Dio2 expression (*Figure 6D*). Although exogenous FGF2 supplementation led to only negligible repression of mitochondrial genes, both Rosi and Lenti-PGC-1α treatment significantly enhanced the mRNA levels of these markers (*Figure 6D*). Furthermore, we observed that Rosi and Lenti-PGC-1α generally acted synergistically (*Figure 6D,E*). It is likely that the FGF2-mediated reduction in UCP1 protein content was counteracted primarily through Rosi activity, while Lenti-PGC-1α played a cooperative role (*Figure 6D,E*). MitoTracker and immunostaining also showed increased accumulation of mitochondria and UCP1 due to Rosi and Lenti-PGC-1α abolition of FGF2-mediated suppression of thermogenic gene expression in brown adipocytes ((*Figure 6F,G*).

In ISO-induced beiging white adipocytes, Rosi largely negated the FGF2 blockade of beiging-related markers, including UCP1, TMEM26, and Tbx1, while Lenti-PGC-1α alone did not significantly restore expression of these markers (*figure supplement 11D*). However, Lenti-PGC-1α substantially antagonized the FGF2 inhibition of mitochondrial marker (CytC, Cox7a, Cox8b, and Aco2) transcription, while Rosi treatment led to only a moderate increase in expression of these markers (*figure supplement 11D*). In addition, the FGF2-reduced UCP1 protein expression was only markedly recovered in treatment of Rosi together with Lenti-PGC-1α (*figure supplement 11E*). In beiging white adipocytes, further evidences for PPARγ and PGC-1α function in the FGF2 pathway for thermogenic gene suppression were observed by Rosi-and Lenti-PGC-1α-restored accumulation of MitoTracker and immunostained UCP1 (*figure supplement 11F,G*).

The expression levels of UCP1 determine thermogenic capability for both brown and beige adipocytes (*Poekes et al., 2015*; *Wu et al., 2012*). Because both PPARγ and PGC-1α function downstream of FGF2 transcriptional suppression of thermogenic genes, we further examined whether FGF2 interfered with interactions between PPARγ/PGC-1α and the UCP1 promoter region. The interaction of PPARγ with PGC-1α was detectable in both brown and beige adipocytes (*Figure 7A,B*). While exogenous FGF2 protein application to cultures of both adipocyte types led to decreased interactions between PGC-1α with PPARγ, the addition of the FGFR1 inhibitor (SSR) alleviated the inhibitory impact of FGF2 (*Figure 7A,B*). Results of chromatin immunoprecipitation (ChIP) assay indicated that PPARγ interaction with the UCP1 promoter was significantly decreased in the presence of FGF2, but largely rescued by Rosi treatment and Lenti-PGC-1α infection. Moreover, Rosi and Lenti-PGC-1α functioned synergistically to recover the FGF2-suppressed UCP1 promoter binding with PPARγ (*Figure 7C,D*). Luciferase reporter activity driven by the UCP1 promoter showed that Rosi and Lenti-PGC-1α both and synergistically restored UCP1 expression that was decreased by FGF2 (*Figure 7E*).

**Fig. 7.**
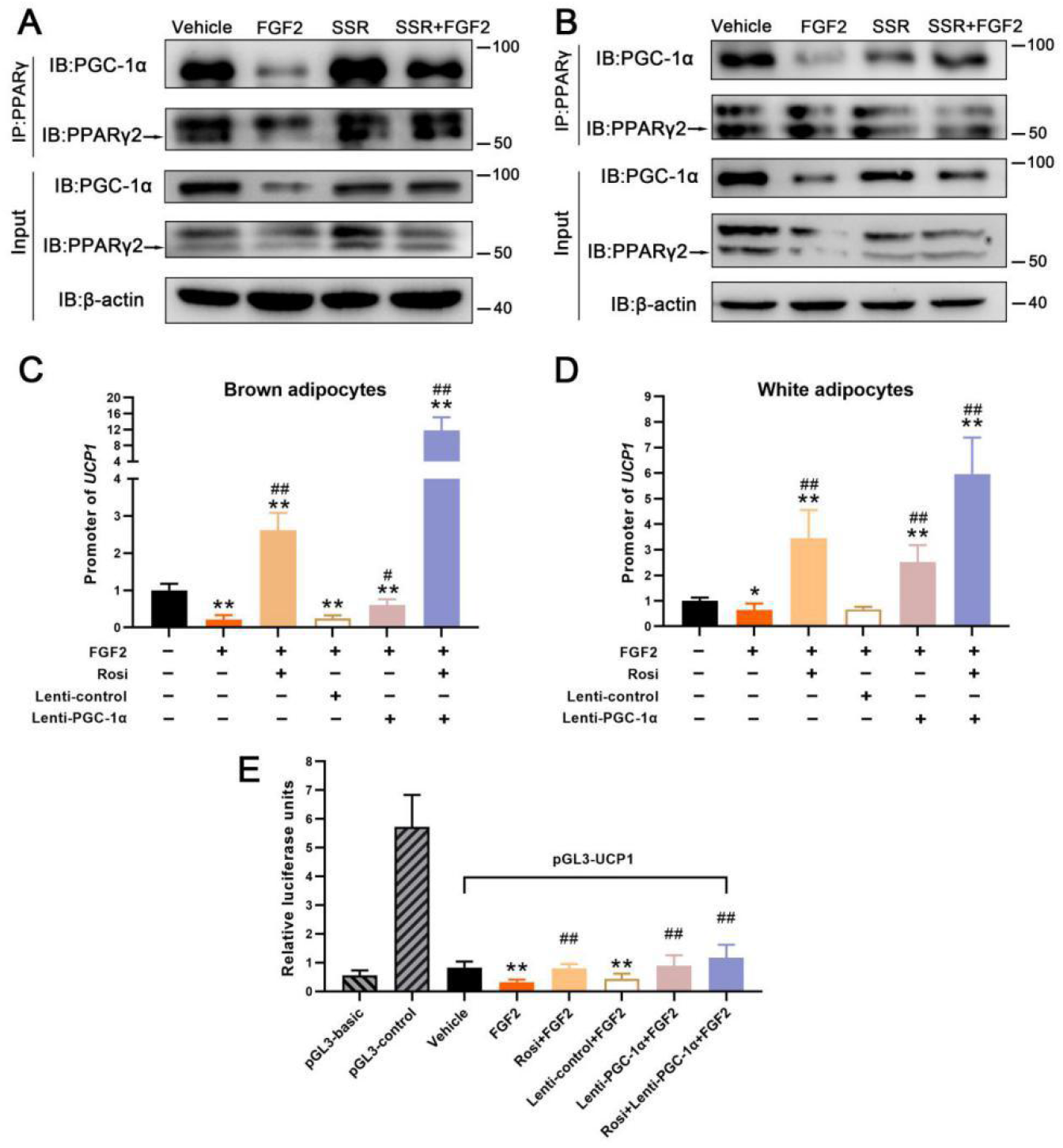
PGC-1α and PPARγ cooperatively participate in FGF2 inhibition of UCP1 expression in both brown and white adipocytes. (*A*) Co-IP analysis of association between PGC-1α and PPARγ in differentiating brown adipocytes treated with Vehicle, FGF2, SSR, or SSR+FGF2. (*B*) Co-IP analysis of association between PGC-1α and PPARγ in differentiating white adipocytes treated with Vehicle, FGF2, SSR, or SSR+FGF2. (*C*) ChIP assay of the association between PPARγ and the UCP1 promoter in brown adipocytes treated with Vehicle, FGF2, Rosi+FGF2, Lenti-control+FGF2, Lenti-PGC-1α+FGF2, or Rosi+Lenti-PGC-1α+FGF2. (*D*) ChIP assay of the interaction between PPARγ and the UCP1 promoter in Vehicle, FGF2, Rosi+FGF2, Lenti-control+FGF2, Lenti-PGC-1α+FGF2, or Rosi+Lenti-PGC-1α+FGF2 treated white adipocytes, in the presence of ISO. (*E*) Dual luciferase activity driven by UCP1 transcription in HEK293 cells treated with Vehicle, FGF2, Rosi+FGF2, Lenti-control+FGF2, Lenti-PGC-1α+FGF2, or Rosi+Lenti-PGC-1α+FGF2. \**p*<0.05, \*\**p*<0.01 vs. Vehicle; ^#^*p*<0.05, ^##^*p*<0.01 vs. FGF2 treatment.

Taken together, these results indicate that although PPARγ and PGC-1α individually provide different contributions in brown and beiging white adipocytes, the two factors work together in the negative regulation of FGF2 on thermogenic gene expression.

## DISCUSSION

Here, we showed that FGF2 disruption stimulated the thermogenic potential of both brown and beige fat, uncovering a previously unrecognized role of FGF2 in adipocyte function. The major findings of this study, as illustrated in *Figure 8*, are as follows: a) FGF2-KO mice exhibit an elevated capacity for thermogenesis in both brown and beige fats, thus showing higher energy expenditure under both basal and β3-AR stimulation; b) FGF2 gene deletion protects against HFD-induced obesity and hepatic steatosis in mice; c) FGF2 leads to ERK phosphorylation, which inhibits the expression of and interactions between PPARγ and PGC-1α, thereby suppressing thermogenic gene expression.

**Fig. 8.**
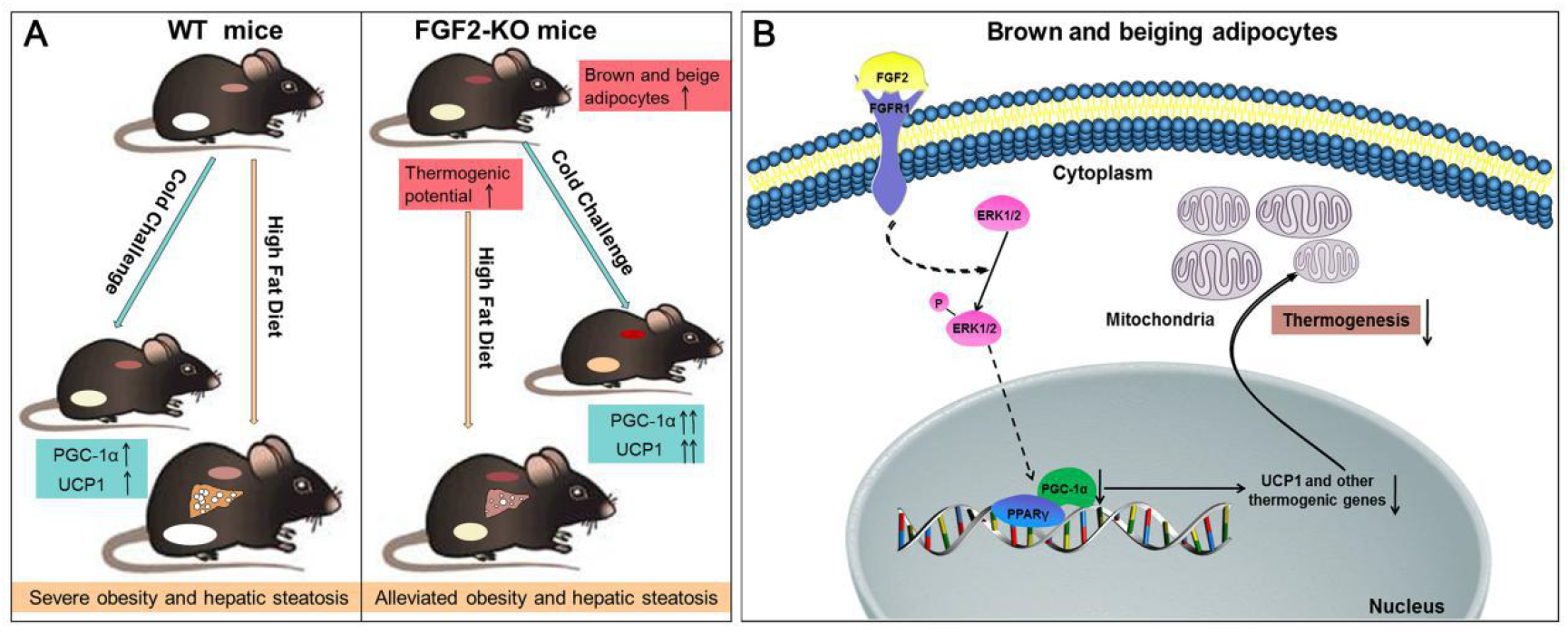
Proposed model of FGF2 regulation the thermogenic potential of brown and beige fat. (*A*) FGF2 deficiency increases brown fat function, and the degree of beiging in white fat, thereby activating thermogenesis under both basal and cold challenge conditions. Consequently, FGF2-KO mice show alleviation of high fat-induced obesity and hepatic steatosis. (*B*) FGF2 stimulates phosphorylation of ERK1/2 in brown and beiging adipocytes and thereafter inhibits the expression of both PPARγ and PGC-1α. The suppression of protein interactions between PPARγ and PGC-1α decreases thermogenic gene expression and the thermogenic potential of brown and beiging adipocytes.

Although several FGF family members, including FGF2, have been established to play unique roles in fat metabolism and/or function, white fat has received considerably more attention (*Jonker et al., 2012*; *Badman et al., 2007*; *Dutchak et al., 2012*; *Sakaue et al., 2002*). For example, Jonker *et al*. reported on the role of the PPARγ-FGF1 signaling axis in adaptive adipose remodeling, in which FGF1-KO mice show impaired adipose expansion under high fat conditions (*Jonker et al., 2012*). FGF21 functions in a feed-forward loop to regulate PPARγ activity via prevention of sumoylation, and FGF21-KO mice show attenuated expression of PPARγ-dependent genes and decreased body fat (*Dutchak et al., 2012*). Furthermore, Fisher *et al*. showed that FGF21 can enhance PGC-1α protein expression as well as the browning of WAT in adaptive thermogenesis in an autocrine/paracrine manner (*Fisher et al., 2012*). Contrary to the role of FGF21, we found that FGF2 negatively affects PGC-1α mRNA and protein abundance, subsequently blocking the white-to-brown fat switch, which was supported by the accumulation of more beige adipocytes in FGF2-KO mice. In addition, we observed an increase in mtDNA copy number and thermogenic gene expression in FGF2-KO BAT. To our knowledge, these findings provide the first description of FGF2 function in thermogenic regulation in both brown and beige fat cells.

As expected, oxygen consumption, RER, and body temperature were all increased due to FGF2 disruption, which was consistent with the increase in whole-body metabolic rate associated with activated brown and/or beige fat function (*Bagchi et al., 2018*; *Yao et al., 2017*). Accordingly, FGF2 disruption amplified the enhancement of cold-induced thermogenic activity for both brown and white fat cells. Thermogenic activity of activated brown and/or beige fat is often accompanied by improved metabolic homeostasis (*Poekes et al., 2015*; *Saito et al., 2009*). Surprisingly, even FGF2^−/−^ mice fed on a chow diet displayed evident improvements in lipid homeostasis. However, adipogenic gene expression increased rather than decreased in KO iWAT and iBAT. This finding supported the potential contribution to improved lipid homeostasis made by increasing thermogenic capability of fat tissues.

In addition, high fat-feeding experiments indicated that FGF2^−/−^ mice were resistant to diet-induced obesity and hepatic steatosis. These results were similar to findings from other studies in which the abundance and/or the thermogenic ability of brown or beige cells were increased in KO strains (such as retinoblastoma protein, tumor necrosis factor-α receptor 1, liver X receptor, and histone deacetylase 11) that also exhibited improved lipid homeostasis and resistance to HFD-induced obesity and/or fatty liver (*Bagchi et al., 2018*; *Hansen et al., 2004*; *Romanatto et al., 2016*; *Wang et al., 2008*). Neither fat synthesis nor fat oxidation was attributable to the triglyceride reduction in the livers of HFD-fed FGF2 KO mice, which further confirmed the contribution of fat thermogenesis on the amelioration of obesity-associated hepatic steatosis phenotypes. These findings suggest a fascinating possibility that priming brown and beige fat function may combat obesity and related metabolic disorders via inhibition of FGF2 signaling.

Given the positive effects of FGF2 disruption on brown fat function and the degree of beiging in white fat cells *in vivo*, we further showed brown and white adipocytes from FGF2-KO mice had higher expression of thermogenic markers *in vitro*, indicating FGF2 took action in an autocrine fashion. Additionally, exogenous FGF2 supplementation suppressed thermogenic gene expression, which demonstrated that FGF2 also acted via paracrine pathways. The cell-autonomous regulation of FGF2 on thermogenesis is similar with that of FGF21, although in contrast, FGF21 enhances the browning of white fat (*Fisher et al., 2012*). Furthermore, we found FGF2 application stimulated ERK1/2 phosphorylation during induced brown adipogenesis and white adipocyte beiging, while blocking ERK signaling counteracted FGF2 suppression of thermogenic genes. These results suggested the involvement of ERK signaling in FGF2-mediated negative regulation of thermogenesis, which is supported by previous reports showing that FGF2 stimulates ERK phosphorylation to inhibit hASC adipogenesis (*Kim et al., 2015*).

We ultimately examined the pathway by which FGF2-induced ERK activation inhibits thermogenic gene expression and discovered that suppression of ERK signaling enhanced both PPARγ and PGC-1α expression, subsequently increasing the abundance of UCP1. PPARγ is a key transcription factor that controls both adipogenic and thermogenic gene expression (*Ahmadian et al., 2013*). However, PPARγ functions coordinately with other components, *e*.*g*., PGC-1α, to activate thermogenesis (*Wu et al., 1999*; *Xue et al., 2005*). Notably, we found that PPARγ and PGC-1α abundance were positively correlated with UCP1 expression *in vivo*. Supplementation with PPARγ-specific agonist and lentiviral expression of PGC-1α *in vitro* blocked FGF2 regulatory function, and synergistically enhanced UCP1 expression, thereby demonstrating the essential regulatory contributions of PPARγ and PGC-1α in modulating FGF2 activity, in agreement with previous studies which showed PGC-1α is a critical transcription co-factors for UCP1 expression (*Wu et al., 1999*; *Xue et al., 2005*). Concurrent with UCP1 activation, PGC-1α expression significantly increased in both FGF2-KO iWAT and iBAT, as well as during cold challenge. Interestingly, Lenti-PGC-1α only moderately increased UCP1 expression, but in the presence of Rosi profoundly elevated UCP1, consistent with previous reports showing that PGC-1α interacts with PPARγ to co-activate genes associated with thermogenesis (*Wu et al., 1999*). In sum, these results indicate that FGF2 application leads to ERK phosphorylation, thereby inhibiting PPARγ and PGC-1α expression and interactions in order to suppress thermogenic gene expression in both brown and beige adipoytes.

In conclusion, we describe an unreported role for FGF2 in the negative regulation of thermogenesis of brown and beige fat in a cell-autonomous manner. FGF2-KO mice show elevated thermogenic ability of brown and beige fat, with higher energy expenditure and improved lipid homeostasis, as well as protection against HFD-induced obesity and hepatic steatosis. Inhibition of PPARγ and PGC-1α expression and interactions via ERK phosphorylation at least partially contributes to the negative regulation of thermogenic gene expression by FGF2. Future studies will further elucidate the contribution of autocrine FGF2 to the inhibition of brown fat function and white-to-beige fat conversion by using mice with adipocyte-specific conditional deletion of FGF2. In addition, further investigation is needed to determine the role of FGF2-specific signaling inhibitors in the thermogenic activities of adipose tissues *in vivo* in order to develop better potential clinical strategies to combat obesity and related disorders.

## Methods

### Reagents

Recombinant FGF2 was from PeproTech (Rocky Hill, USA). PD and Rosi were purchased from Selleck (USA). CL was from Cayman Chemical (USA). PGC-1α expression lentivirus (Lenti-PGC-1α) and control lentivirus (Lenti-control) were from Genechem (Shanghai, China). The following primary antibodies were used: GAPDH (AB0038, Abways, China), β-actin (P60709, Abways, China), FGF2 (sc-74412, Santa Cruz), PPARγ (sc-7273, Santa Cruz), PGC-1α (sc-13067, Santa Cruz), UCP1 (ab15517, Abcam), aP2 (sc-271529, Santa Cruz), ERK (#4695, CST), and p-ERK (#9101, CST).

### Animals

All experiments were conducted in FGF2-KO (FGF2^−/−^) mice and WT (FGF2^+/+^) littermates with the same genetic background (C57BL/6J). KO was conducted using clustered regularly interspaced short palindromic repeats (CRISPR)/CRISPR-associated 9 methods. Two single-guide RNAs (sgRNA1: GGAGACAGAGGCCTGCAATG and sgRNA2: TCTCGCGGACGCCATCCAC G) were designed to target promoter and exon 1. Successful deletion was confirmed by PCR genotyping using tail genomic DNA with primers 5’-TCTAACAACTGAGGCAGGGCAA-3’ and 5’-GAAGTGGCAACTCAC CGTG TG-3’. FGF2 heterozygous (+/–) mice were bred to obtain FGF2-KO mice and their WT littermates.

Mice were housed in a temperature- and humidity-controlled, pathogen free facility with 12 h dark-light cycles. For HFD studies, animals were fed a diet that 45% kcal from fat (Ref. D12451, Research Diets Inc., USA). The body weight, food and water intake of mice were recorded weekly. The body composition of mice was determined by dual energy X-ray absorptiometry (DXA) (MEDIKORS, Korea). At the end of the experiment, animals were kept fasting for 12 h and sacrificed by isoflurane inhalation followed by cervical dislocation. iBAT, iWAT, eWAT and liver tissues were harvested and weighed. All animal experiments were performed in accordance with the guidelines for the Care and Use of Laboratory Animals, and animal maintenance and experimental procedures were approved by the Animal Care and Use Committee of Shandong Agricultural University, China.

### Histology, H&E staining, ORO staining, and immunohistochemistry/ immunofluorescence analysis

iBAT, iWAT, eWAT and liver tissues were fixed in 4% paraformaldehyde for more than 24 h, and embedded in paraffin. Paraffin samples were sectioned (5 μm) and stained with hematoxylin and eosin for histochemical examination. Area of adipocytes in adipose tissues were measured by using an image software (Nikon, Japan). For ORO staining, liver samples were frozen in liquid nitrogen and sectioned at 8 μm in thickness using a cryostat. The sections were stained with ORO solution for 10 min. After washing with water, sections were stained for 1 min in hematoxylin. For immunohistochemistry/immunofluorescence, de-paraffinized iBAT and iWAT sections were blocked with FBS, incubated with specific UCP1 primary and HRP-/Alexa Fluor 555-conjugated secondary antibodies, and detected accordingly.

### Scanning electron microscopy (SEM)

iBAT and iWAT samples were fixed with 2% glutaraldehyde, and post-fixed in 1% osmium tetroxide for 1 h, dehydrated in graded concentrations of ethanol and 100% acetone. The specimens were dried at the critical point. Subsequently, the specimens were stuck on a colloidal silver, and sputtered with gold by a MED 010 coater (Balzers) and analyzed with a scanning electron microscope (JEOL, Japan).

### Quantification of mtDNA copy number

Equal amounts of WT and KO iBAT and iWAT were used to extract total DNA after digestion with proteinase K, respectively. The isolated DNA was used to amplify mtDNA using primers for the mitochondrial cytochrome c oxidase subunit 2 (COX2) gene, with the Rsp18 nuclear gene as an internal control of genomic DNA, as described previously (*Yao et al., 2017*).

### Determination of plasma parameters

Whole blood was collected from eyeball into heparinized containers and plasma was obtained after centrifugation. Fasting triglyceride (TG) and total cholesterol (TCH) levels were determined by commercial kits (Njjc Bio Institute, China). Plasma alanine aminotransferase (ALT) and aspartate aminotransferase (AST) activities were measured on Cobas Integra 400 Clinical Analyzer (Roche Diagnostics).

### Hepatic TG and TCH content determination

Liver tissue (500 mg) was homogenized in 300 μL RIPA lysis buffer in a Polytron disrupter. The homogenate was centrifuged at 12,000g for 5 min, and the supernatant was collected. TG and TCH content in the tissue was quantified with commercial assay kits (Dongou, China), which was normalized to total protein and expressed as mmol/g total protein.

### Metabolic studies

Mice were housed in individual metabolic cage system (TSE LabMaster, TSE system, Germany) with free access to water and food, and allowed to acclimate for a 24 h period, then data was collected every 9 min for another 24 h. O_2_ consumption (VO_2_) and CO_2_ production (VCO_2_) were measured by the TSE system, and RER were calculated using the manufacturer’s system software.

### Infrared thermography

The temperature of WT and KO mice was recorded with an infrared camera (FOTRIC 225) and analyzed with a specific software (FOTRIC Tools Software). At least five pictures of each mouse were taken and analyzed.

### Cold challenge and β3-AR agonist treatment

For cold exposure experiment, WT and FGF2^−/−^ mice were kept at a 5 °C room for 24 hours. Core body temperature was monitored using a rectal probe every 3 or 6 hours for the 24 h-duration of the study. Twenty four hours’ later, mice at RT and cold room were sacrificed to collect iBAT and iWAT for further thermogenic-capability determinations. For β3-AR agonist treatment, CL was intraperitoneally injected into mice at 0.5 mg kg^−1^ body weight/day. Three days later, iBAT and iWAT were collected to determine the mRNA expression of thermogenic genes and make paraffin slices.

### Isolation of brown and white SVFs, in *vitro* differentiation and treatments

Brown and white SVFs were obtained and induced to differentiate into mature adipocytes, respectively, as previously described with minor modification (*Seale et al., 2011*; *Bagchi et al., 2018*). During the induced adipogenic process, 1 μM PD, 1 μM Rosi or 0.5 μM SSR were added to some cell culture dishes, in the presence or absence of 10 ng/mL FGF2 and 10 ng/mL HP.

### Quantitative real-time PCR (qRT-PCR)

Total RNA was extracted from tissues or cells using RNAiso Plus Reagent, and converted to cDNA using the HiScript ? Q RT SuperMix for qPCR Kit (Vazyme, China). qRT-PCR was performed with SYBR green fluorescent dye (Takara, Japan) using a Real-Time PCR System (Applied Biosystems). Transcript levels were quantified using the 2^-ΔΔCt^ method values to that of GAPDH. Primers used were shown in *table supplement 1*.

### Western blotting

Protein lysates were obtained from tissues or cells using RIPA lysis buffer containing protease and phosphatase inhibitor cocktails (Roche, USA). Western blotting was conducted as described previously (*Yao et al., 2017*).

### Lentivirus infection

For experiments with lentivirus, different recombinant lentiviruses were individually supplemented into 50% confluency SVFs at 5 MOI with 6 μg/mL polybrene, and the medium was refreshed 24 hours after infection. After recovering for another 72 hours, the infected cells were induced for differentiation together with other treatments.

### MitoTracker staining

Different-treated brown and beige adipocytes were stained with MitoTracker Red CMXRos (20 nM) (CST, #9082) in DMEM containing 15% FBS at 37 °C for 30 min. Following washing twice with DMEM containing 15% FBS, the cells were incubated with DAPI (1 μg/mL) for 5 min at RT. The intracellular MitoTracker-stained mitochondria were detected using a confocal laser scanning microscopy (CLSM) (Zeiss, Germany). Images were acquired and processed with the same setting for different treatments.

### Immunofluoresence staining of UCP1

For immunofluorescence staining, formalin-fixed and Triton X 100-permeabilized brown and beige adipocytes were pre-incubated with a blocking buffer (PBS containing 5% FBS) for 60 min, and incubated with UCP1 antibody (1:100 dilution) in blocking buffer at 4 °C overnight. Subsequently, the slides were washed, and incubated with Alexa Flour 555-conjugated secondary antibody (1:500 dilution). After staining with DAPI (1 μg/mL) for 5 min, images were acquired by using a CLSM (Zeiss, Germany) and processed with the same setting for different treatments.

### Co-IP

To determine the influence of FGF2 and/or FGFR1 inhibitor SSR on interaction of PPARγ with PGC-1α in differentiating brown and beige adipocytes, co-IP was performed as previously described (*Bagchi et al., 2018*), with some minor modifications.

### ChIP assay

To determine the interaction of PPARγ with the promoter of UCP1, CHIP assay was performed using a CHIP Assay Kit, as described previously (*Yao et al., 2017*). The primers for UCP1-promoter were listed in *table supplement 1*.

### Dual luciferase reporter assay

HEK293 cells cultured in 96-well plates were cotransfected with 1000 ng/mL of pGL3-basic, pGL3-UCP1p, or pGL3-control along with 50 ng/mL of pRL-TK, respectively. Six hours later, the cells were infected with Lenti-control or Lenti-PGC-1α recombinant viruses in some treatments. After infection with 24 hours, the cells were treated with Vehicle, FGF2, Rosi+FGF2, Lenti-control+FGF2, Lenti-PGC-1α+FGF2, or Rosi+Lenti-PGC-1α+FGF2 for 2 days. Subsequently, the treated cells were washed and lysed in 20 μL of lysis buffer (Dual reporter assay system, Promega). The firefly luciferase activity was examined according to the protocols, and efficiency was normalized to renilla luciferase activity directed by a cotransfected control plasmid pRL-TK.

### Statistical analysis

Statistical analysis was performed on data from at least 3 repeated experiments. All data were presented as means±SEM. Significant difference between treatments was tested by one-way ANOVA or two-sample student t-test. p<0.05 was regarded as significant.

## Acknowledgments

The authors would like to thank Zizhang Zhou (Shandong Agricultural University), Xuguo Zhou (University of Kentucky), and Yingli Shang (Shandong Agricultural University) for their assistance in preparation of this manuscript. Thanks will be given to Yanqin Wang (Hebei Normal University) for assistance with lentivirus production. This work was supported by the Taishan Scholars Program [No. 201511023], the National Key Research and Development Program of China (2017YFE0129800), Funds of Shandong “Double Tops” Program, and Natural Science Foundation of Shandong Province, China [No. ZR2019MC016].

## Notes

### Competing Interest Statement

The authors have declared no competing interest.

