## Supplementary material for "FGF2 disruption enhances thermogenesis in brown and beige fat to protect against obesity and hepatic steatosis": figure supplement 1

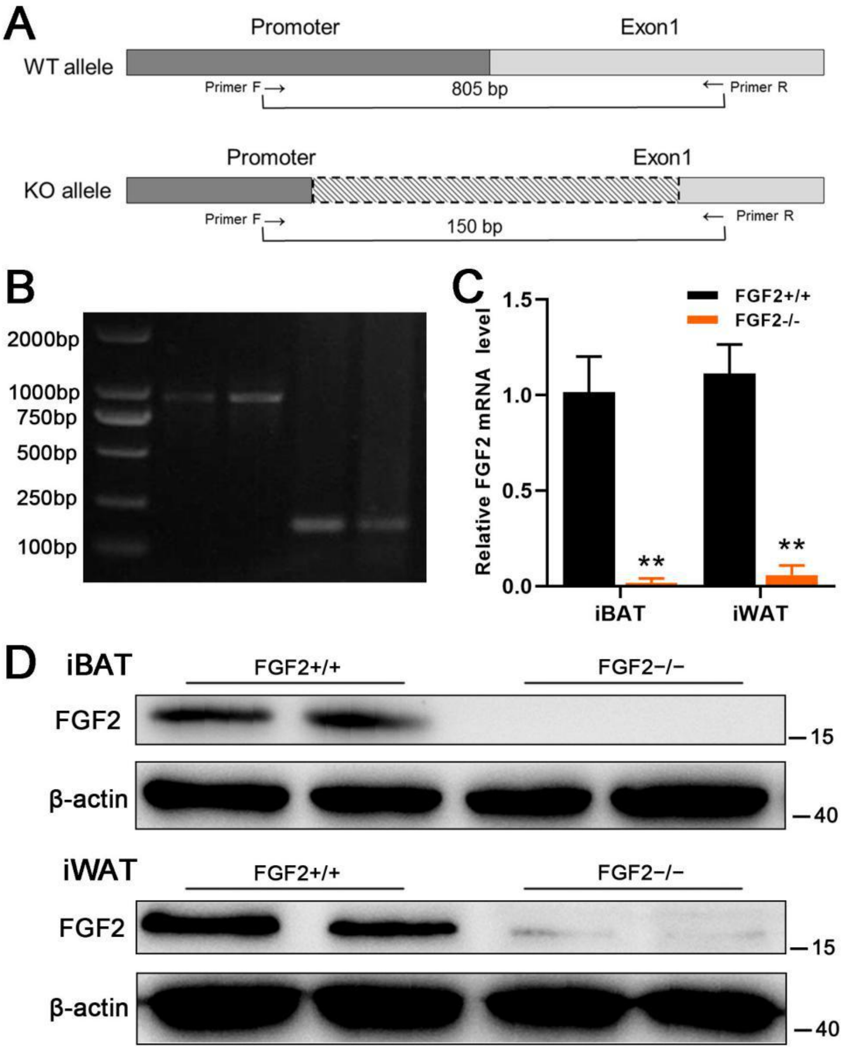


**Fig. S1. Preparation and identification of FGF2-KO mice.**

(A) Shown (top to bottom) were wild-type and deleted FGF2 gene loci. Primer pairs were for genotyping were designed to distinguish WT and KO alleles, and sizes of the expected PCR products are indicated. (B) Genotyping results of $FGF2^{+/+}$ and $FGF2^{-/-}$ mice. (C) Relative mRNA levels of FGF2 in iBAT and iWAT from $FGF2^{+/+}$ and $FGF2^{-/-}$ mice. Values for fold induction of gene expression were normalized to GAPDH and expressed as means ± SEM (n = 6). *p<0.05, **p<0.01 for $FGF2^{-/-}$ versus $FGF2^{+/+}$. (D) Western blot analysis of FGF2 protein expression in iBAT and iWAT from $FGF2^{+/+}$ and $FGF2^{-/-}$ mice. β-actin serves as a loading control.
