## Supplementary material for "FGF2 disruption enhances thermogenesis in brown and beige fat to protect against obesity and hepatic steatosis": figure supplement 2

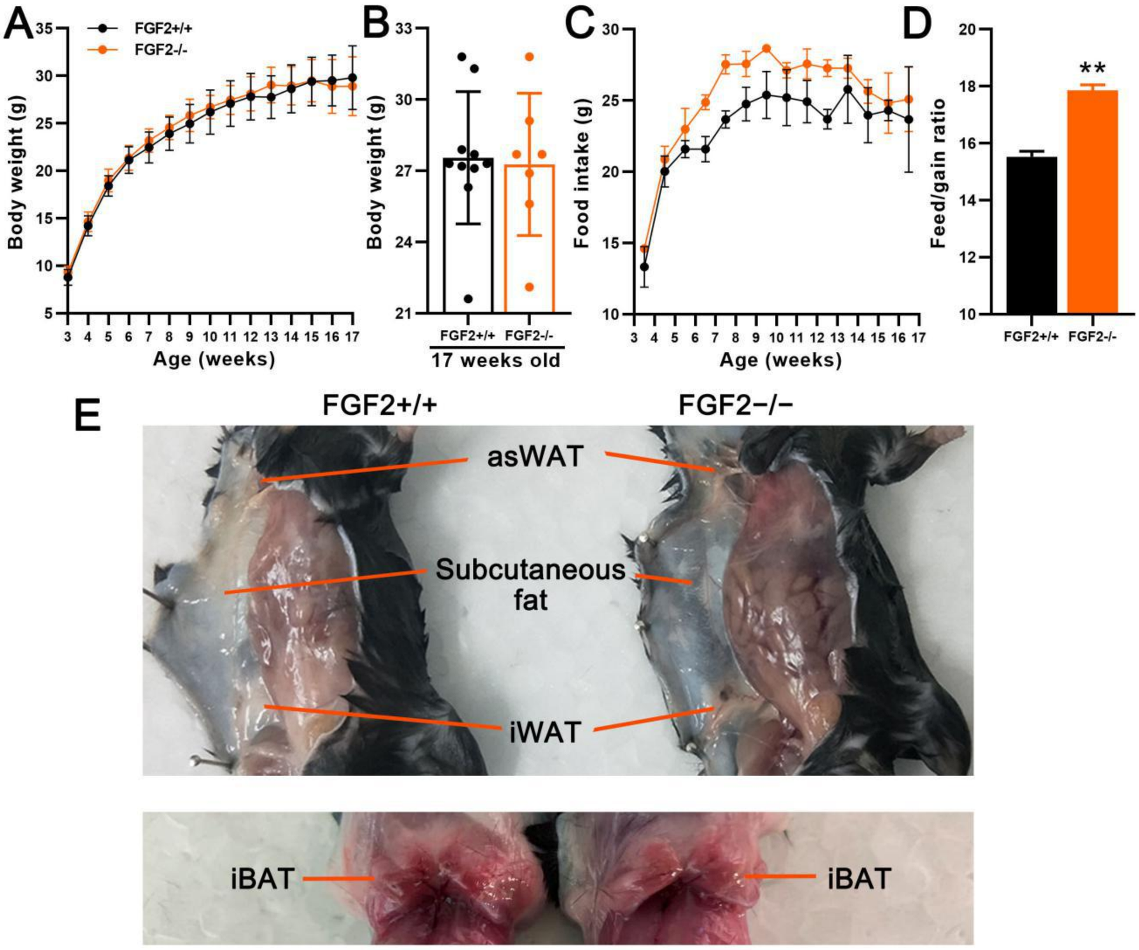


**Fig. S2. Comparison of body weight, food intake, and anatomy images in FGF2$^{+/+}$ mice with those of FGF2$^{–/–}$ mice.**

3-week-old male FGF2$^{+/+}$ and FGF2$^{–/–}$ mice were fed with chow diet for 14 weeks, and then euthanized at the age of 17 weeks. (A) Dynamic changes in body weight of FGF2$^{+/+}$ and FGF2$^{–/–}$ mice during over the full experimental time course. (B) Body weights of 17-week-old mice. (C) Dynamic shifts in intake of FGF2$^{+/+}$ and FGF2$^{–/–}$ mice over the course of the whole expressed showing as g/mouse/week. (D) The whole feed/gain ratio of FGF2$^{+/+}$ and FGF2$^{–/–}$ mice for 14 weeks. (E) Representative anatomical images of 17-week-old mice. Values represent means ± SEM (n = 7~10). **p<0.01 compared with FGF2$^{+/+}$ mice.
