## Supplementary material for "FGF2 disruption enhances thermogenesis in brown and beige fat to protect against obesity and hepatic steatosis": figure supplement 3

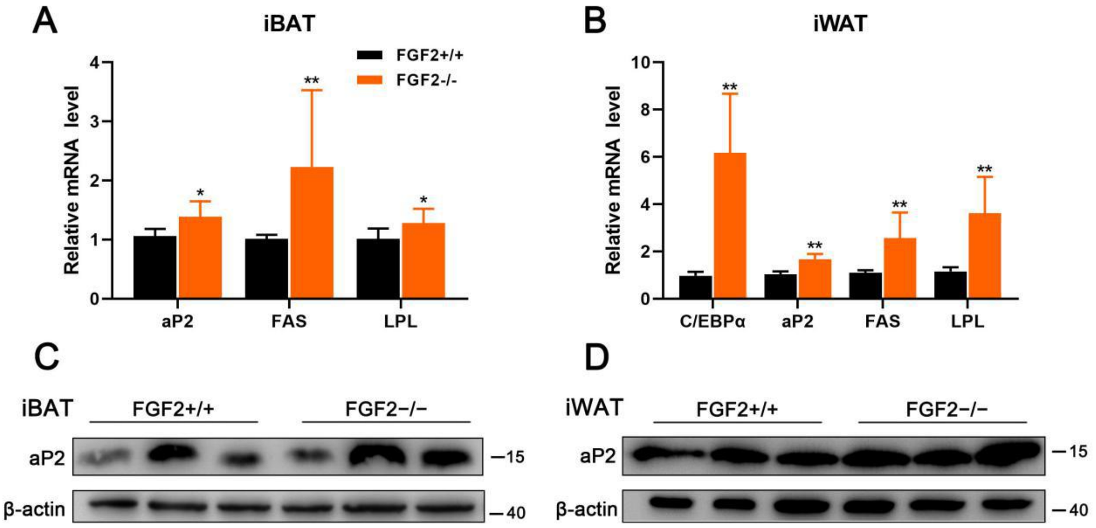


**Fig. S3. Adipogenic gene expression levels were enhanced in iBAT and iWAT of FGF2$^{-/-}$ mice, compared with those of FGF2$^{+/+}$ mice.**

Tissue samples were from 17-week-old male FGF2$^{+/+}$ and FGF2$^{-/-}$ mice. (A and B) The relative mRNA levels of adipogenic markers in iBAT (A) and iWAT (B) of FGF2$^{+/+}$ and FGF2$^{-/-}$ mice determined by qRT-PCR. Values represent means ± SEM (n = 6). *p<0.05, **p<0.01 compared with FGF2$^{+/+}$ samples. (C and D) Western blot analysis of the protein content of aP2 in iBAT (C) and iWAT (D) of FGF2$^{+/+}$ and FGF2$^{-/-}$ mice. Blots were stripped and probed again with β-actin to normalize for variation in loading and transfer of proteins.
