## Supplementary material for "FGF2 disruption enhances thermogenesis in brown and beige fat to protect against obesity and hepatic steatosis": figure supplement 4

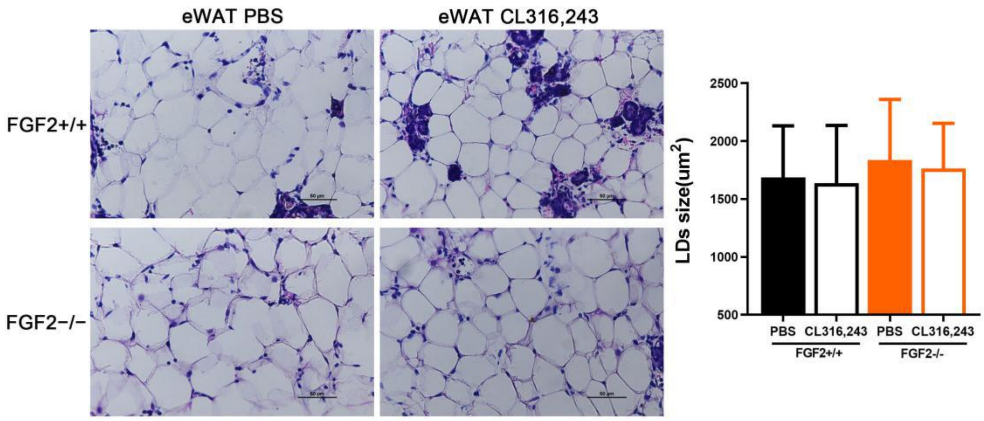


**Fig. S4. eWAT is unresponsive to CL316,243 injection.**

Representative images of H&E staining of eWAT tissue sections and the lipid droplet sizes, upon vehicle (PBS) or CL316,243 injection. Scale bar = 50 μm.
