## Supplementary material for "FGF2 disruption enhances thermogenesis in brown and beige fat to protect against obesity and hepatic steatosis": figure supplement 5

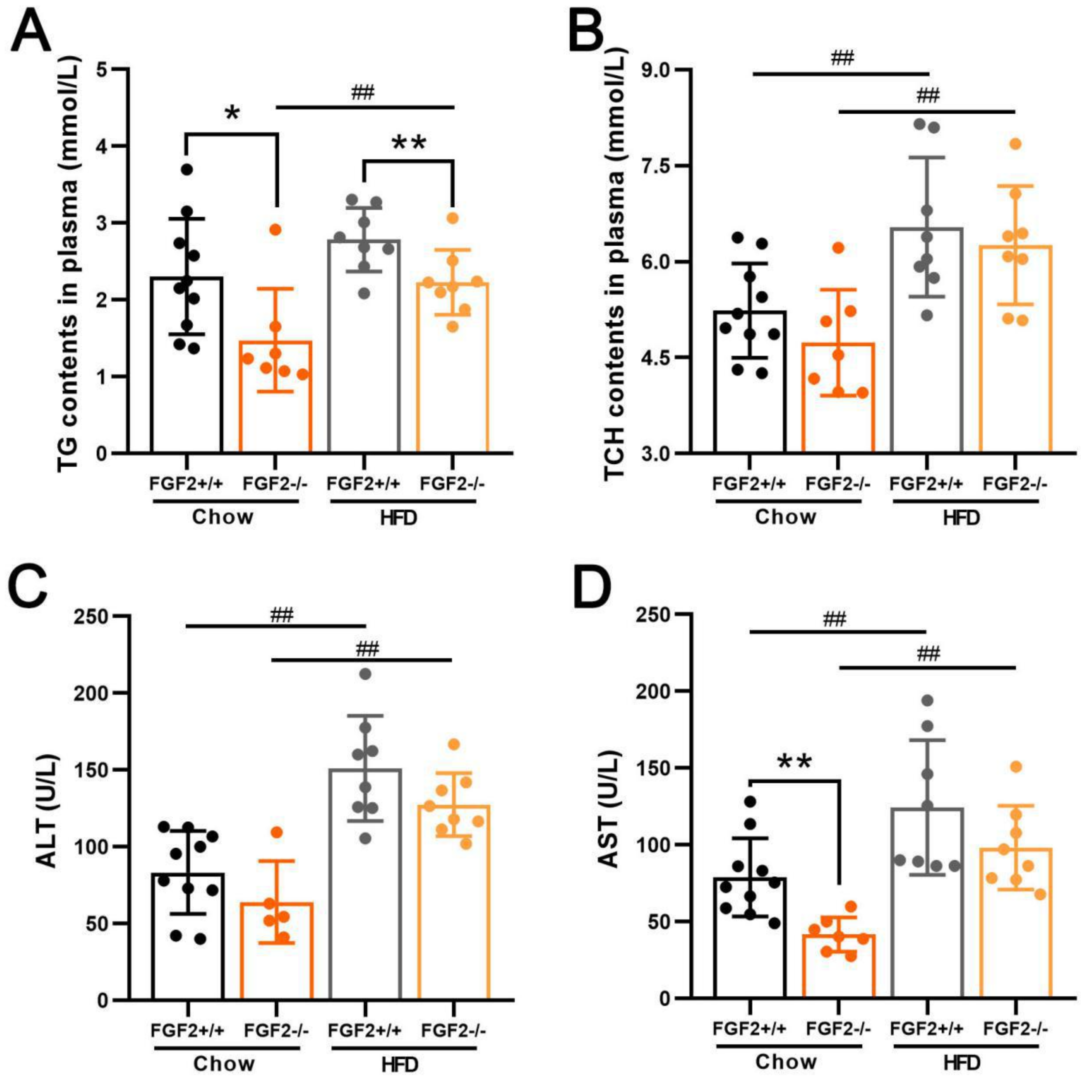


**Fig. S5. FGF2$^{-/-}$ mice show improved lipid homeostasis.**

Samples were from 17-week-old FGF2$^{+/+}$ and FGF2$^{-/-}$ mice fed with chow or high fat diet. (A-D) Plasma TG (A) and TCH (B) contents, as well as ALT (C) and AST (D) activities in FGF2$^{+/+}$ and FGF2$^{-/-}$ mice (chow diet and HFD). Values represent means ± SEM (n = 7~10). *p<0.05, **p<0.01 compared with FGF2$^{+/+}$ mice feeding with the same diet; $^{\#}$p<0.05, $^{\#\#}$p<0.01 compared with the same genotype fed with chow diet.
