## Supplementary material for "FGF2 disruption enhances thermogenesis in brown and beige fat to protect against obesity and hepatic steatosis": figure supplement 6

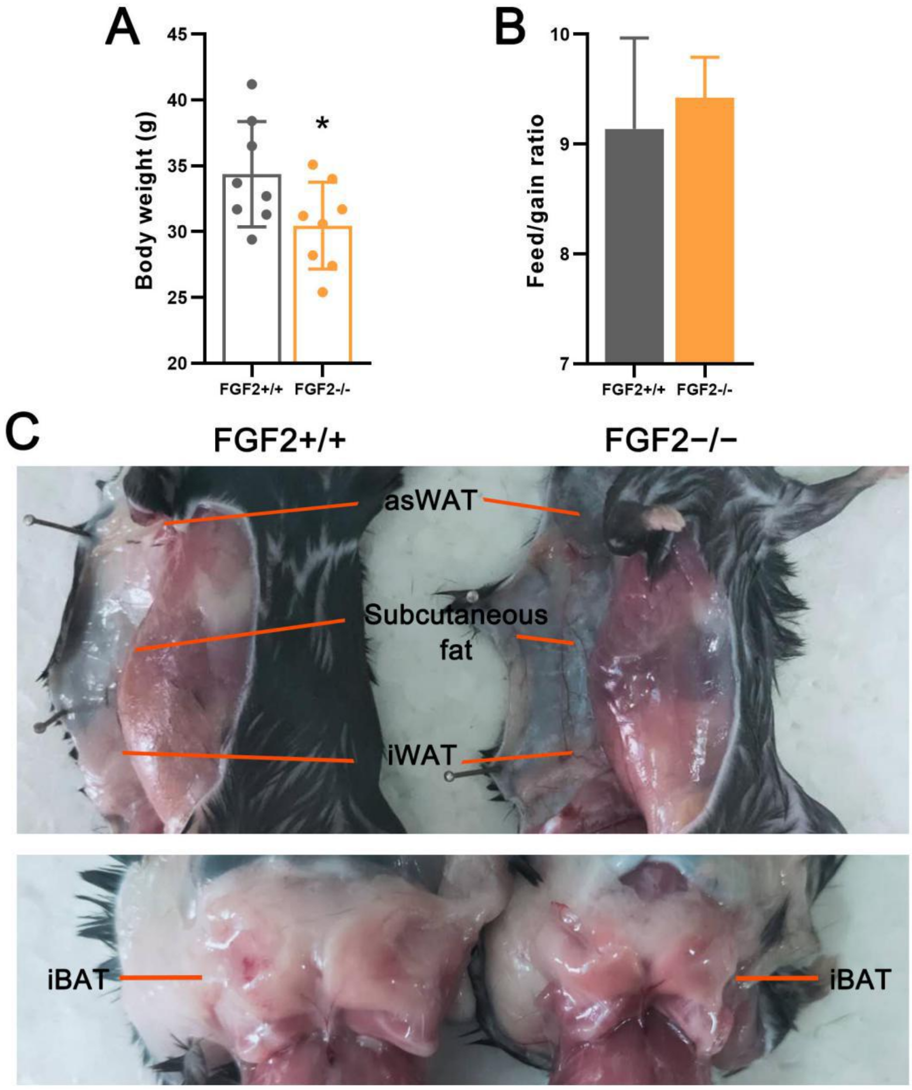


**Fig. S6. The body weight, food intake, and anatomical images of $FGF2^{+/+}$ and $FGF2^{-/-}$ mice upon HFD feeding.**

3-week-old male $FGF2^{+/+}$ and $FGF2^{-/-}$ mice were fed with HFD for 14 weeks and then euthanized at the age of 17 weeks. (A) Body weight of 17-week-old $FGF2^{+/+}$ and $FGF2^{-/-}$ mice fed with HFD. (B) The whole feed/gain ratio of $FGF2^{+/+}$ and $FGF2^{-/-}$ mice for 14 weeks. (C) Representative anatomical images of 17-week-old $FGF2^{+/+}$ and $FGF2^{-/-}$ mice fed with HFD. Values represent means ± SEM (n = 8). *p<0.05, **p<0.01 compared with $FGF2^{+/+}$ mice.
