## Supplementary material for "FGF2 disruption enhances thermogenesis in brown and beige fat to protect against obesity and hepatic steatosis": figure supplement 7

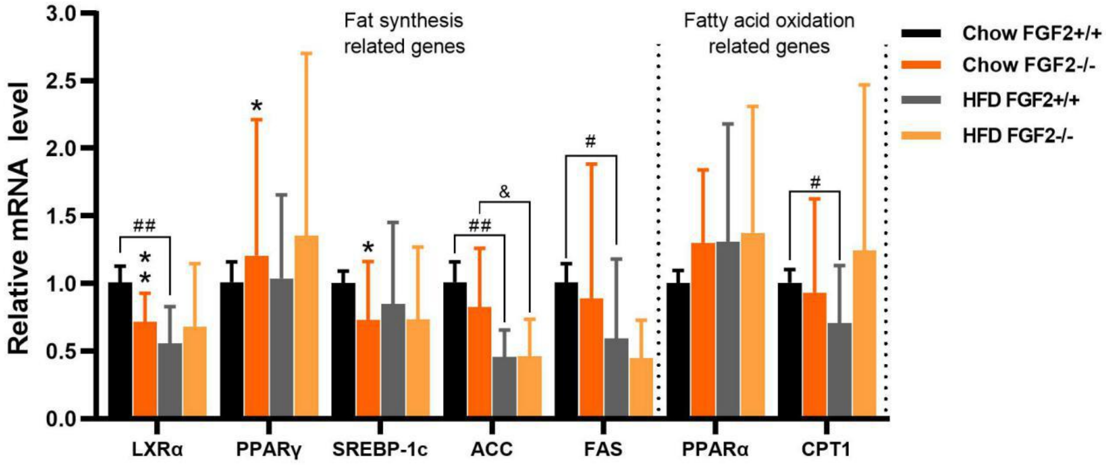


**Fig. S7. Fat synthesis- and lipolysis-related gene expression in livers of FGF2$^{+/+}$ and FGF2$^{–/–}$ mice fed with chow or high fat diet.**

Data were from qRT-PCR analysis, using GAPDH as a reference. Data represent means ± SEM (n = 6).*p<0.05, **p<0.01 vs. FGF2$^{+/+}$ mice fed with the same diet; $^{\#}$p<0.05, $^{\#\#}$p<0.01 vs. FGF2$^{+/+}$ mice fed with chow diet; $^{\&}$p<0.05 vs. FGF2$^{–/–}$ mice fed with chow diet.
