## Supplementary material for "FGF2 disruption enhances thermogenesis in brown and beige fat to protect against obesity and hepatic steatosis": figure supplement 8

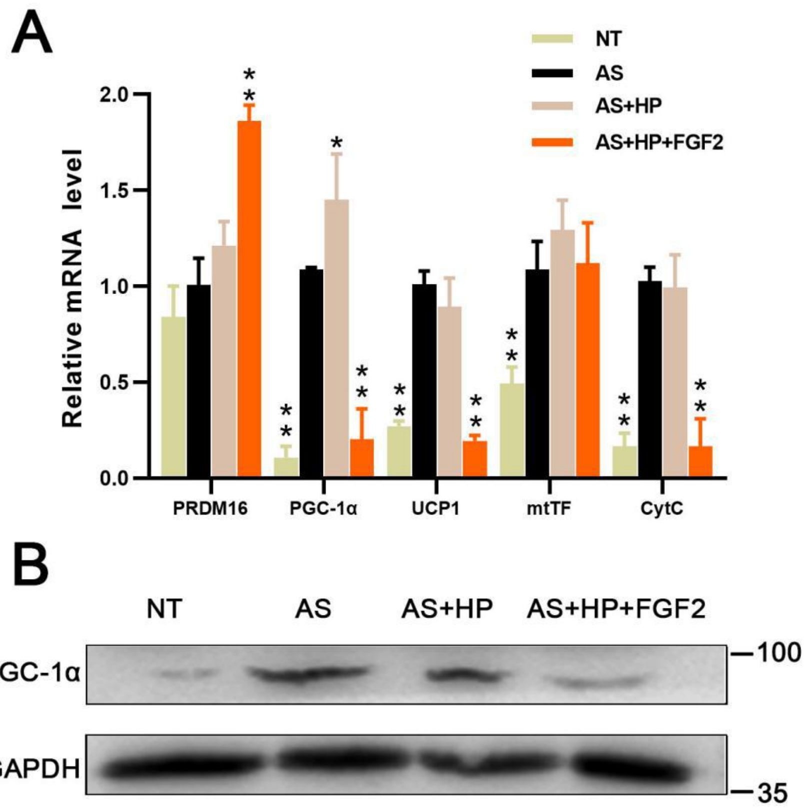


**Fig. S8. FGF2 inhibits thermogenic gene expression in brown adipocytes in vitro.**

(A) Relative mRNA expression of genes involved in brown adipocyte function (PRDM16, PGC-1α, UCP-1, mtTF and Cytc) in cells treated with NT (no treatment), AS (adipogenic stimuli), AS+HP (10 ng/mL), or AS+HP (10 ng/mL)+FGF2 (10 ng/mL). (B) Western blot analysis of PGC-1α protein expression in cells with each indicated treatment. GAPDH served as a loading control. Data represent means ± SEM.*$p<0.05$, **$p<0.01$ compared with AS treatment.
