## Supplementary material for "FGF2 disruption enhances thermogenesis in brown and beige fat to protect against obesity and hepatic steatosis": figure supplement 9

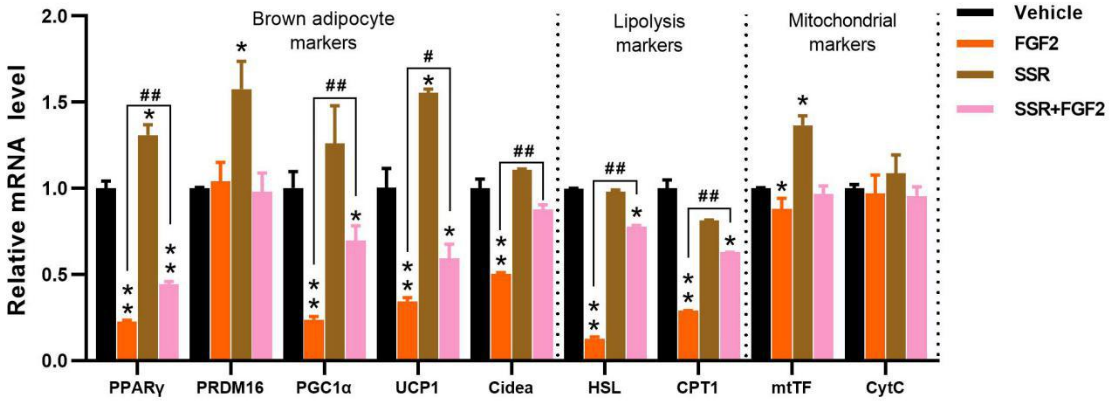


**Fig. S9. Exogenous FGF2 application inhibits thermogenic gene expression in brown adipocytes in a paracrine manner.**

The inhibition of FGF2 on thermogenic genes expression could be partially rescued by SSR (a FGFR1 inhibitor), as indicated by qRT-PCR analysis. GAPDH serves as an internal control. Data represent means ± SEM (n = 6).$^{*}p<0.05$, $^{**}p<0.01$ vs. Vehicle; $^{\#}p<0.05$, $^{\#\#}p<0.01$ vs. FGF2 treatment.
