## Supplementary material for "FGF2 disruption enhances thermogenesis in brown and beige fat to protect against obesity and hepatic steatosis": figure supplement 10

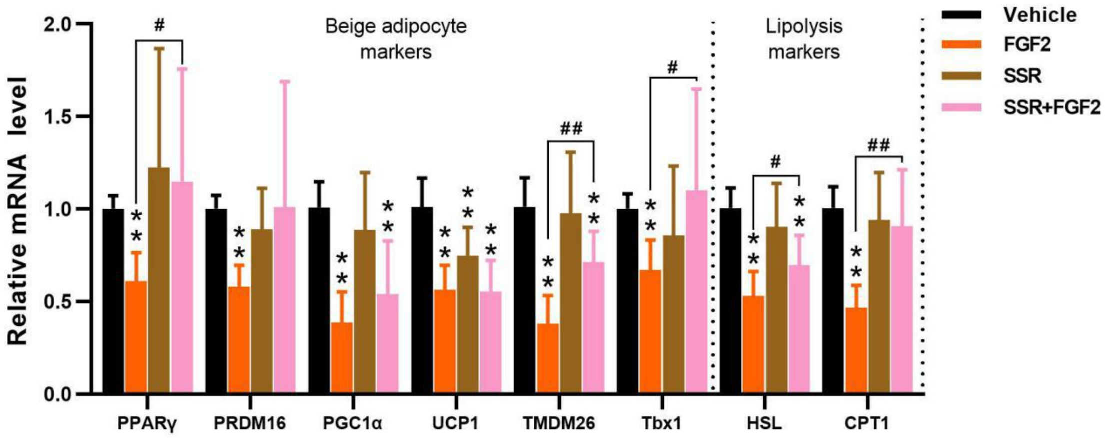


**Fig. S10. FGF2 affects on beiging-associated gene expression in white adipocytes in a paracrine fashion.**

The suppression of FGF2 on beiging-associated gene expression could be rescued by SSR, as indicated by qRT-PCR analysis of samples treated with vehicle, FGF2, SSR, or SSR+FGF2. GAPDH served as an internal control. Data represent means ± SEM (n = 6). *p<0.05, **p<0.01 vs. Vehicle; $^{\#}$p<0.05, $^{\#\#}$p<0.01 vs. FGF2 treatment.
