## Supplementary material for "FGF2 disruption enhances thermogenesis in brown and beige fat to protect against obesity and hepatic steatosis": figure supplement 11

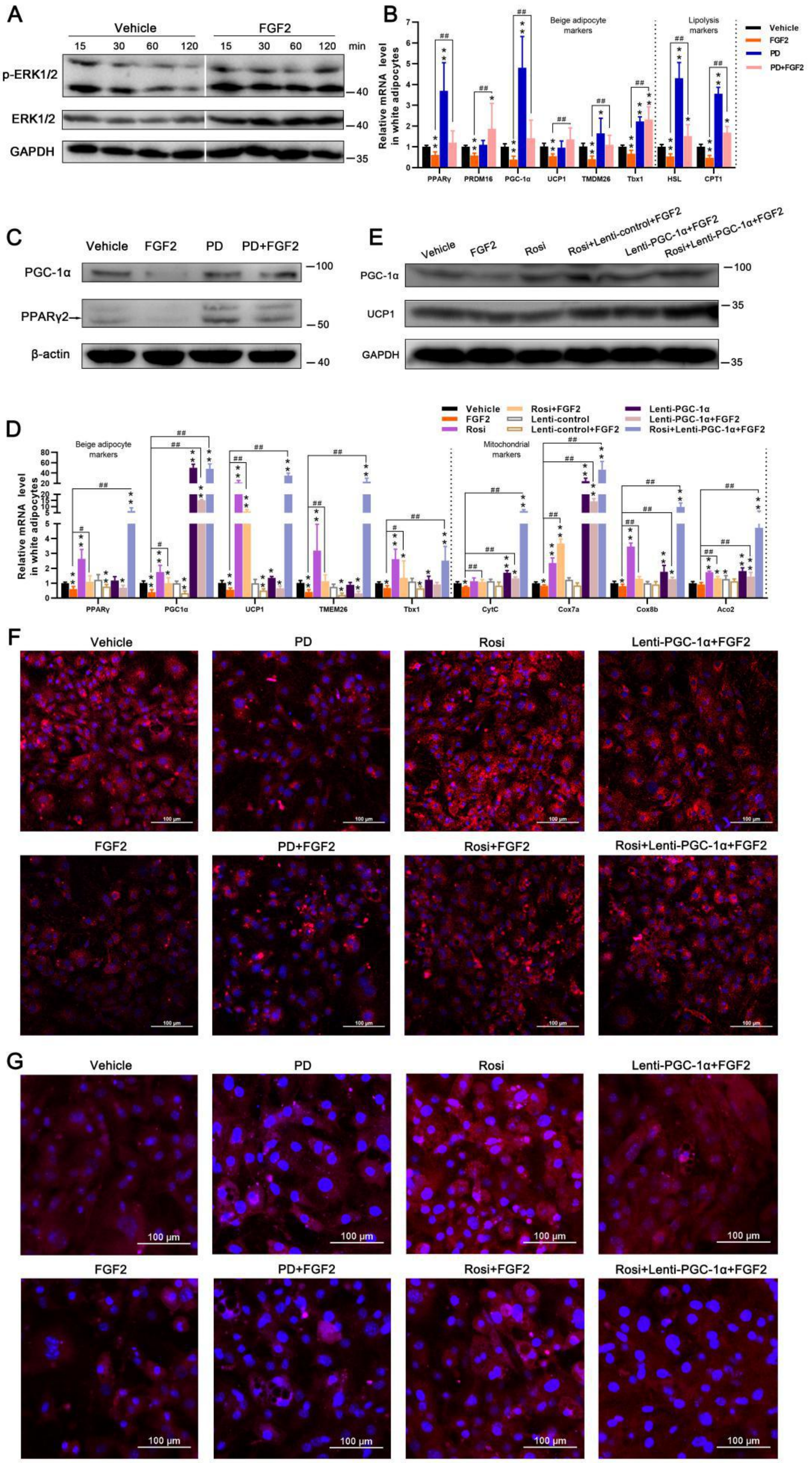
A
Vehicle
FGF2
15 30 60 120 15 30 60 120 min
p-ERK1/2
ERK1/2
GAPDH
B
Relative mRNA level in white adipocytes
Beige adipocyte markers
Lipolysis markers
Vehicle
FGF2
PD
PD+FGF2
PPARγ PRDM16 PGC-1α UCP1 TMDM26 Tbx1 HSL CPT1
C
Vehicle FGF2 PD PD+FGF2
PGC-1α
PPARγ2
β-actin
E
Vehicle FGF2 Rosi Rosi+Lenti-control+FGF2 Lenti-PGC-1α+FGF2 Rosi+Lenti-PGC-1α+FGF2
PGC-1α
UCP1
GAPDH
D
Vehicle
FGF2
Rosi
Rosi+FGF2
Lenti-control
Lenti-control+FGF2
Lenti-PGC-1α
Lenti-PGC-1α+FGF2
Rosi+Lenti-PGC-1α+FGF2
Beige adipocyte markers
Mitochondrial markers
Relative mRNA level in white adipocytes
PPARγ PGC1α UCP1 TMEM26 Tbx1 CytC Cox7a Cox8b Aco2
F
Vehicle PD Rosi Lenti-PGC-1α+FGF2
FGF2 PD+FGF2 Rosi+FGF2 Rosi+Lenti-PGC-1α+FGF2
G
Vehicle PD Rosi Lenti-PGC-1α+FGF2
FGF2 PD+FGF2 Rosi+FGF2 Rosi+Lenti-PGC-1α+FGF2
100 μm

**Figure S11. FGF2 suppresses ISO-initiated thermogenic gene expression in white adipocytes *in vitro*, via ERK signaling-induced PPARγ and PGC-1α inhibition.**

(A) Expression of ERK1/2 and p-ERK1/2 proteins in differentiating white SVFs after supplementing with FGF2 or Vehicle for 15, 30, 60, and 120 min, determined by western blotting. (B) Relative mRNA levels of beige adipocyte- and lipolysis-associated genes in Vehicle, FGF2, PD, or PD+FGF2 -treated white adipocytes, in the presence of ISO. GAPDH serves as a loadling control. (C) Protein expression of PPARγ, and PGC-1α in white adipocytes treated as in (B), determined by western blotting. β-actin serves as a loading control. (D) Relative mRNA levels of beige adipocyte- and mitochondrial-associated genes in Vehicle, FGF2, Rosi, Rosi+FGF2, Lenti-control, Lenti-control+FGF2, Lenti-PGC-1α, Lenti-PGC-1α+FGF2, or Rosi+Lenti-PGC-1α+FGF2 -treated white adipocytes, in the presence of ISO. (E) Protein expression of PGC-1α and UCP1 in ISO-incubated white adipocytes treated with Vehicle, FGF2, Rosi, Rosi+FGF2, Lenti-PGC-1α+FGF2, or Rosi+Lenti-PGC-1α+FGF2, determined by western blotting. (F and G) MitoTracker staining (red) (F) and immunofluorescence of UCP1 (red) (G) of treated white adipocytes, in the presence of ISO. The nuclei (blue) were stained with DAPI. Scale bar = 100 μm. Data represent means ± SEM. *$p<0.05$, **$p<0.01$ vs. Vehicle; $^{\#}p<0.05$, $^{\#\#}p<0.01$ vs. FGF2 treatment.
