## Supplementary material for "FGF2 disruption enhances thermogenesis in brown and beige fat to protect against obesity and hepatic steatosis": table supplement 1

**Table S1. The primers used in this study.**

| Genes | Sequence 5' to 3' |
|---|---|
| GAPDH | Forward-AGAGTGTTTCCTCGTCCCG<br>Reverse-CCGTTGAATTTGCCGTGA |
| FGF2 | Forward-TCCCACCAGGCCACTT<br>Reverse-CGTCCATCTTCCTTCATA |
| Primer-GT | Forward-TCTAACAACTGAGGCAGGGCAA<br>Reverse-GAAGTGGCAACTCACCGTGTG |
| PPARγ | Forward-AGACACCCTTTCACCAGCATCC<br>Reverse-AACCCTTACAACCTTCACAAGCA |
| aP2 | Forward-CCTTTGTGGGAACCTGGAA<br>Reverse-TGTCGTCTGCGGTGATTT |
| C/EBPα | Forward-TCGGTGCGTCTAAGATGAGG<br>Reverse-TGAGTATCCAAGGCACAAGGT |
| FAS | Forward-GGGTCTATGCCACGATTC<br>Reverse-GTGTCCCATGTTGGATTTG |
| LPL | Forward-GAGGATGGCAAGCAACAC<br>Reverse-AGCAGTTCTCCGATGTCC |
| SREBP-1c | Forward- GACTACATCCGCTTCTTGC<br>Reverse- CACCACTTCGGGTTTCAT |
| ACC | Forward- AGGCTATGTGAAGGATGTG<br>Reverse- CTGAAGAGGTTAGGGAAG |
| PPARα | Forward- CAAGTGCCTGTCTGTCGG<br>Reverse- GCGGGTTGTTGCTGGTCT |
| HSL | Forward-AGCACTACAAACGCAACGA<br>Reverse-CGACAGCACCTCAATCTCA |
| CPT1 | Forward-CGTGACGTTGGACGAATC<br>Reverse-TCTGCGTTTATGCCTATC |
| PGC-1α | Forward-AGAAGCGGGAGTCTGAAA<br>Reverse-CAGGTGTAACGGTAGGTG |
| PRDM16 | Forward-GCGGTCAGCAATAGCAGC<br>Reverse-CCCGTGGTAGTGTCCAAGTC |
| UCP1 | Forward-GCTTAATGACTGGAGGTGTG<br>Reverse-GCTTTCTGTGGTGGCTAT |
| Cidea | Forward-CTTCCTCGGCTGTCTCAA<br>Reverse-TGGCTGCTCTTCTGTATCG |

| | |
|---|---|
| C/EBPβ | Forward-TGTCCACGTCGTCGTCGTCC<br>Reverse-CCGTCAGCTCCAGCACCTTGT |
| TMEM26 | Forward-GACTCCACCAAACACTCC<br>Reverse-GCATACTCCACGTCCACA |
| Tbx1 | Forward-ACCGAGATGATCGTCACCAAG<br>Reverse-ACCAGCCAGGAGGAGCTATG |
| mtTF | Forward-CCTCGTCTATCAGTCTTGT<br>Reverse-GCTTCTGGTAGCTCCCTC |
| CytC | Forward-ATCTCCACGGTCTGTTCG<br>Reverse-GCCCTTTCTCCCTTCTTC |
| Cox7a | Forward-GGCTCTGGTCCGGTCTT<br>Reverse-CTGGGAGGTCATTGTCG |
| Cox8b | Forward-TGCGAAGTTCACAGTGGTT<br>Reverse-GGCGGAAGTGGGAGTTTT |
| Aco2 | Forward-ATCGAGCGGGGAAAGACATAC<br>Reverse-TGATGGTACAGCCACCTTAGG |
| Uqcrc2 | Forward-AAAGTTGCCCCGAAGGTTAAA<br>Reverse-GAGCATAGTTTTCCAGAGAAGCA |
| COX2 (mtDNA) | Forward-ATAACCGAGTCGTTCTGCCAAT<br>Reverse-TTTCAGAGCATTGGCCATAGAA |
| Rsp18 (Genomic DNA) | Forward-CATCACCCACTTACCCCCAAAA<br>Reverse-TGTGTTAGGGGACTGGTGGACA |
